# Wounding perception by jasmonate-mediated JAZ condensation in plants

**DOI:** 10.1101/2024.03.28.587122

**Authors:** Zi-Wei Yan, Jie Liu, Hui-Cheng Chen, Mu-Yang Wang, Wen-Juan Cai, Jian-Xu Li, Jia-Wei Wang, Ying-Bo Mao

## Abstract

Wounding is a common stress experienced by plants, resulting in a quick elicitation of jasmonates (JAs). The widely accepted mechanism of COI1-JAZ coreceptor posits that COI1 directly interacts with the active ligand of JAs, whereas JAZs facilitate to trap the ligand in the binding pocket. Here we show that the multiple JAZs of *Arabidopsis* undergo phase separation in response to wounding. Taking JAZ1 as an example, we found it is an independent sensor of JA-Ile, the active ligand in most plants. Under basal condition, dispersed JAZ1 interacts with MYC2 to inhibit downstream gene expressions. Upon wounding, JAZ1 binds to JA-Ile and forms condensates, reducing its affinity with MYC2 and tending to recruit COI1 as well as other proteasome related components for quick degradation, thereby accelerating signaling output. Our investigation unveils the underlying mechanism by which JAZ1 regulates swift signaling transduction via phase separation upon the direct perception of JA-Ile.

## Introduction

As sessile organisms, plants inevitably encounter wounding arising from various abiotic and biotic stresses like insect herbivores^1,2^. The lipid-derived phytohormone jasmonates (JAs) are the predominant regulators in plant wounding response^3,4^. In the absence of external stimulus, the contents of JAs are maintained at a low level in plants^5^. JASMONATE ZIM-domain (JAZ) proteins which are the main repressors inhibit JA signaling transduction by binding to the transcription factors like MYC2^6^. A quintuple mutant of JAZs (*jaz1 jaz3 jaz4 jaz9 jaz10*, *jazQ*) displays a hypersensitive response to JA and wounding treatment^7^. Wounding triggers an outbreak of JAs in plants^8^. JA-Ile, the active ligand in most plants, directly interacts with CORONATINE INSENSITIVE1 (COI1), an integral component of the SKP1-CUL1-F-box protein (SCF) E3 ubiquitin ligase complex known as SCF^COI19^, and triggers the interaction between COI1 and JAZs, leading to the degradation of JAZs through the 26S proteasome^10^. The crystal structure has been identified that the Jas motif of *Arabidopsis* JAZ1 contributes to trap JA-Ile in the binding pocket of COI1^11^. Furthermore, JAZs also assist COI1 in determining ligand specificity^12^. However, in contrast to COI1, no evidence of the independent recognition and interaction of active JAs with JAZs has been reported so far^13,14^. It is commonly accepted that COI1 primarily perceives active JA ligand and this process is assisted by JAZs.

It has been discovered that many kinds of proteins form liquid-like droplets or compartments *in vivo*, which is required for their functions^15–17^. In plants, phase separation has been found to be involved in diverse biological processes^18–21^. NPR1, the master regulator of salicylic acid signaling, become condensed to improve cell survival upon the plant immune response^21^. Phase separation of the nuclear pore complex promotes the nucleocytoplasmic transport of immune regulators, thereby enhancing anti-insect defense in plants^22^. Proteins capable of phase separation usually harbor the intrinsically disordered region (IDR)^23^. Rather than a fixed, rigid structure, IDR is distinguished by its inherently flexible and diverse conformational assembles^24,25^, which provides multivalent interactions required for driving condensation. Some IDR included proteins have been identified as sensors which detect and respond to various environmental stimuli through the dynamic phase separation^17,26,27^. TERMINATING FLOWER (TMF) in plant shoot apical meristem, undergoes condensation upon the perception of reactive oxygen species (ROS) to maintain the synchronizing flowering^28^. In plant interspecific competitions, RNA-BINDING PROTEIN 47B (RBP47B) perceives donor plant-secreted phenolic acids (PAs) via phase separation, thereby leading to the formation of stress granules and translational inhibitions in neighbor plants^29^.

Here, we found multiple JAZs in *Arabidopsis* harbor IDRs and undergo phase separation in the perception of wounding. Taking JAZ1 for example, we found that the wounding-triggered JAZ1 condensation depends on JA-Ile rather than COI1. MST analysis reveals an interaction between JAZ1 and JA-Ile. And the *in vitro* condensation of JAZ1 can be directly promoted by the additional JA-Ile. In dispersed phase, JAZ1 shows higher binding affinity with MYC2, whereas in condensed phase, JAZ1 is more inclined to interact with COI1 and recruits other proteasome related components for quick degradation. The capacity of JAZ1 switch from dispersed to condensed phase allows a prompt and sensitive signaling transduction. Our investigation unveils that JAZs act as direct sensors to perceive elicited JA-Ile upon wounding via phase separation.

## Results

### JAZ proteins undergo phase separation in plants upon wounding

The *Arabidopsis* genome encodes a total of 13 JAZ proteins, which can be classified into five distinct clades based on phylogenetic analyses (Extent Data Fig. 1a). Except for JAZ7, JAZ8, JAZ9 and JAZ11, the remaining JAZ proteins harbor intrinsically disordered region (IDR), as predicted by the PONDR^30^ program (Extent Data Fig. 1b). Besides, IDRs are also present in JAZ proteins from *Marchantia polymorpha* (liverwort), *Huperzia selago* (lycophyte), *Oryza sativa* (monocot) plants (Extent Data Fig. 2) reflecting a conserved evolution. From the reported transcriptome data of *Arabidopsis*^31^, the expression of most JAZs can be significantly induced by wounding whereas JAZ4 and JAZ13 are hardly detected in leaf tissues (Extent Data Fig. 3). The JAZ proteins containing IDR and showing detectable transcription levels in leaf tissues were fused with Venus (V-JAZs), a green fluorescent protein, and transiently expressed in *N. benthamiana* leaves to investigate their capability of phase separation *in vivo*. V-JAZs are located in nucleus as accepted and we noticed near the edge of the cut-out leaf discs (Zone 2), which are close to the wounded side, all the 7 tested JAZ proteins are in condensed phase whereas at Zone 1 far away from wounded site, they are in dispersed phase (Fig. 1a and Extent Data Fig. 4a). We proposed that JAZs might undergo phase separation, which is driven by wounding. To test this hypothesis, laser wounding was carried on the pavement cells in Zone1 followed by single-cell time-lapse imaging. Taking JAZ1 as an example, it rapidly undergoes condensation after laser wounding, and the number of condensates per cell shows a gradual increase within two minutes (Fig. 1b, c and Supplementary Video 1). In addition, we observed that two approaching condensates fused into a bigger one (Fig. 1d and Supplementary Video 2), and the fluorescent intensity within condensates recovered after photobleaching (Extent Data Fig. 4b, c and Supplementary Video 3). These results suggest that the JAZ1 condensates are driven by liquid-liquid phase separation upon wounding.

**Fig. 1:**
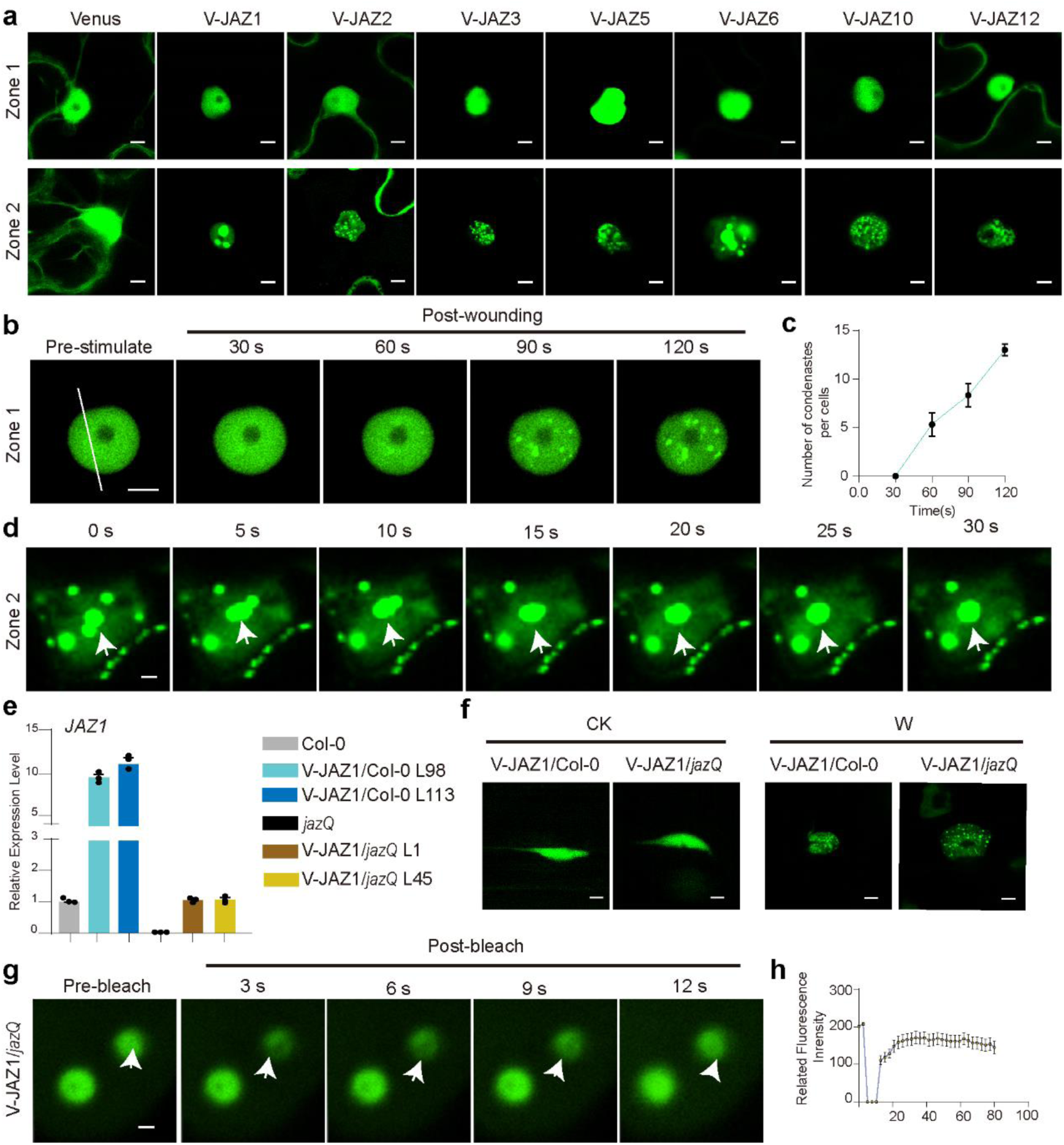
Wounding triggers the condensation of multiple JAZs in plants. **a**, JAZ proteins form condensates within Zone 2 (around the wounding sites) rather than Zone 1 (far away from the wounding sites) (See also Extended Data Fig. 3a). JAZ proteins with IDR were fused with Venus. Venus and Venus-fused JAZ proteins (V-JAZs) are transiently expressed in the leaves of *N. benthamiana*. Leaf discs are observed under confocal microscopy. Scale bars, 5 μm. **b,** V-JAZ1 at Zone 1 rapidly undergoes condensation upon laser wounding. The imaging process is initiated immediately after wounding (time 0) and lasted for 2 minutes. Images from indicated time points are selected. Scale bars, 5 μm. **c,** Quantification of JAZ1 condensates per cell at the indicated time points following laser wounding as described in (**b**). Data are mean ± SEM (n = 3 biological replicates). **d,** Fusion events of V-JAZ1 condensates at Zone 2. The arrow indicates the fusion location. Scale bar, 1 μm. **e,** qRT-PCR analysis of *JAZ1* expression in Col-0, *jazQ* and transgenic plants. *JAZ1* expression level in *35S:V-JAZ1 jazQ* (V-JAZ1/*jazQ*) is parallel to those in wild type (Col-0). Data are mean ± SEM (n = 3 biological replicates). **f,** JAZ1 condensates were observed in both *35S:V-JAZ1* line 98 (V-JAZ1/Col-0) and *35S:V-JAZ1 jazQ* line 1 (V-JAZ1/*jazQ*) upon wounding. The untreated (CK) and wounded (W) roots were observed under confocal microscopy. Scale bars, 5 μm. **g,** Fluorescence recovery after photobleaching (FRAP) of JAZ1 condensates in the wounded roots of *35S:V-JAZ1 jazQ* line 1 (V-JAZ1/*jazQ*). The arrow indicates the bleached condensate. Scale bars, 1 μm. **h,** Quantification of the recovered fluorescence intensity after photobleaching as described in (**g**). Data are mean ± SEM (n = 6 biological replicates).

V-JAZ1 was then introduced into *Arabidopsis* plants of the wild-type Col-0 and the JA-hypersensitive mutant *jazQ*^7^ driven by the 35S promoter (*35S:V-JAZ1* and 35S: *V-JAZ1 jazQ*). The transcription levels of *JAZ1* in *35S:V-JAZ1* (line 98 and 113) were more than 10 times higher than those in Col-0, whereas *35S:V-JAZ1 jazQ* (line 1 and 45) showed a comparable transcription level of *JAZ1* as Col-0 (Fig. 1e). In immune blot assay, the JAZ1 levels in transgenic plants are consistent to their transcription level (Extent Data Fig. 5a). Similar to those observed in *N. benthamiana*, the distribution of JAZ1 is dispersed throughout the nucleus in both untreated plants of *35S:V-JAZ1* and *35S:V-JAZ1 jazQ* while it coalesces into condensates in response to wounding (Fig. 1f). When the *35S:Venus* plants were used for investigation, no condensation was observed regardless of wounding (Extent Data Fig. 5b). And the comparable transcription level of *JAZ1* in *35S:V-JAZ1 jazQ* and Col-0 (Fig. 1e) excludes the possibility that the condensation is caused by an overdose. From the spatiotemporal analysis of bleaching events, the fluorescence intensity of JAZ1 condensates in the roots of *35S:V-JAZ1* (Extent Data Fig. 5b,d and Supplementary Video 4) and *35S:V-JAZ1 jazQ* (Fig. 1g, h and Supplementary Video 5) rapidly restored after photo bleaching indicating that JAZ1 condensates are highly dynamic. These results suggest that JAZ1 undergoes liquid-liquid phase separation in the perception of wounding in *Arabidopsis*.

### JA is required for wounding-triggered JAZ1 condensation

It is well known that wounding triggers a burst of JAs in most plants^1^. To investigate whether JAs are involved in JAZ1 condensation upon wounding, we treated leaf discs of *N. benthamiana* transiently expressing V-JAZ1 with methyl-JA (MeJA) which activates the JA signaling pathway^32^. The dispersed V-JAZ1 in Zone 1 gradually transits into condensates in the presence of MeJA (Fig. 2a, b and Supplementary Video 6) which is very similar to that of laser-wounding treatment (Fig. 1b). Consistently, JAZ1 undergoes condensation in *35S:V-JAZ1* seedlings upon both wounding and MeJA treatments (Fig. 2C and Extent Data Fig. 6b), whereas Venus along is unable to form condensates (Extent Data Fig. 6a). We next introduced V-JAZ1 into *aos* (*35S:V-JAZ1 aos*), a mutant deficient in JA synthesis^33^. Two transgenic lines (2 and 9), of which the JAZ1 expressions were examined by both qRT-PCR and immune blot assay (Extent Data Fig. 7a, b), were selected for investigation. We found that JAZ1 condensates are only observed in *35S:V-JAZ1 aos* upon MeJA but not wounding treatment (Fig. 2d and Extent Data Fig. 7c). Together these results indicate that successful JA synthesis is essential for wounding-triggered JAZ1 condensation in plants.

**Fig. 2:**
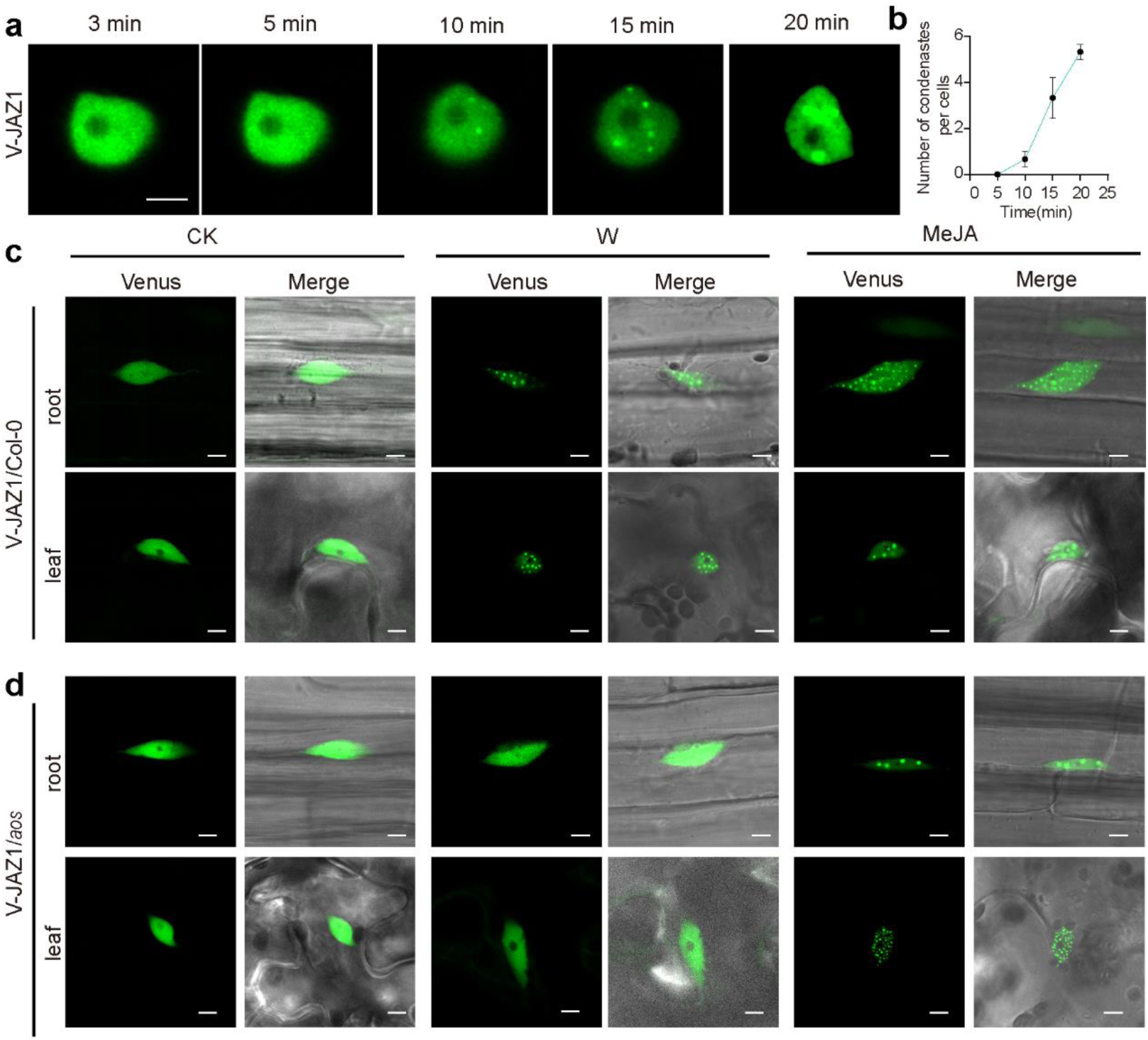
Wounding-mediated JAZ1 condensates are dependent on JA. **a**, Time-lapse imaging of V-JAZ1 in *N. benthamiana* post JA treatment. Images from Zone 1 are started after 1 mM MeJA addition and lasted for 20 min. Scale bars, 5 μm. **b,** Quantification of JAZ1 condensates per cell post MeJA treatment as described in (**a**). Data are mean ± SEM (n = 3 biological replicates). **c-d,** Observations of V-JAZ1 in *Arabidopsis* plants. CK represents the unwounded plants. W represents the wounded plants. MeJA represents the plants treated with 50 μM MeJA. The pavement cells of the leaf and elongation region cells of the roots from the plants of *35S:V-JAZ1* line 98 (V-JAZ1/Col-0) and *35S:V-JAZ1 aos* line 2 (V-JAZ1/*aos*) were observed under confocal microscopy immediately post treatment. Scale bars, 5 μm.

### JAZ1 is a direct sensor of JA-Ile independent of COI1

So far, only COI has been reported to directly perceive the active ligand of JAs^13,14,34^. To evaluate whether the formation of JAZ1 condensation depends on COI1, we next expressed V-JAZ1 in *coi1-2*^35^ (*35S:V-JAZ1 coi1-2*). The mRNA and protein levels of JAZ1 in transgenic plants (line 2 and 4) were examined (Extent Data Fig. 8a, b). V-JAZ1 in the root and leaf cells of *35S:V-JAZ1 coi1-2* transgenic plants are unable to form condensates upon MeJA treatment, however, both wounding and JA-Ile do induce its condensate formations (Fig. 3a and Extent Data Fig. 8c). We then detected the levels of JA and JA-Ile in *35S:V-JAZ1 coi1-2* plants after MeJA or wounding treatments. It reveals that MeJA treatment leads to a higher accumulation of JA rather than JA-Ile, whereas wounding elicits the accumulation of JA-Ile instead of JA (Fig. 3b, c). This reflects a robust positive correlation between JA-Ile and JAZ1 condensation. The prokaryotically expressed mCherry-JAZ1, not mCherry, spontaneously forms droplets which positively correlate with the protein concentration (Extent Data Fig. 9a-d). The droplet formation is inhibited by 1,6-hexanediol due to its disruption of hydrophobic interaction, while was not disturbed by NaCl (Extent Data Fig. 9e,f). The *in vitro* JAZ1 condensates can also recover fluorescent intensity after photobleaching (Extent Data Fig. 9g, h and Supplementary Video 7), and fuse together (Extent Data Fig. 9i and Supplementary Video 8), suggesting a comparable physical nature to that of *in vivo* (Fig. 1 d, g).

**Fig. 3:**
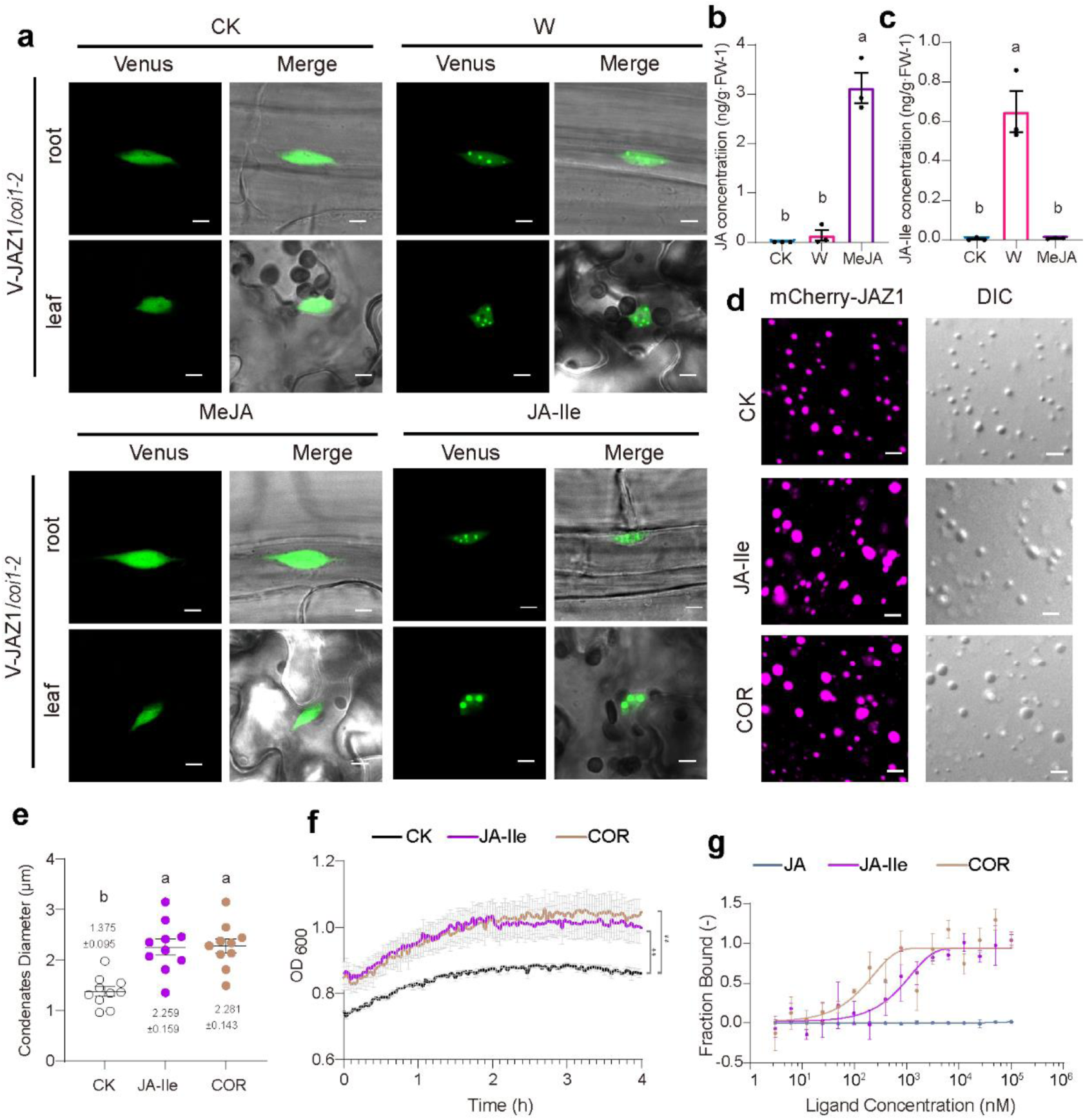
JAZ1 directly binds to JA-Ile and undergoes phase separation in a COI1-independent manner. **a,** The JAZ1 condensation in *Arabidopsis* is dependent on JA-Ile rather than on COI1. The plants of *35S:V-JAZ1 coi1-2* line 2 (V-JAZ1/*coi1-2* were wounded (W), and treated with 50 μm of MeJA (MeJA) or JA-Ile (JA-Ile). Pavement cells of the leaf and elongation region cells of the root were immediately observed under confocal microscopy. CK represents untreated plants. Scale bars, 5 μm. **b-c,** Quantification of JA (**b**) and JA-Ile (**c**) concentrations in *coi1-2* mutants as described in (**a**). Data are analyzed by one-way ANOVA followed by two-sided Tukey’s multiple comparisons test. Different letters represent significant differences (*P* < 0.05). Data are mean ± SEM (n = 3 biological replicates). **d,** The JA-Ile and Coronatine directly promote the JAZ1 phase separation *in vitro*. About 10 μM JAZ1 without additives (CK), and with 1 mM of JA-Ile or Coronatine (COR) were observed under confocal microscopy. DIC indicates differential interference contrast. Scale bars, 5 μm. **e,** The diameter measurement of JAZ1 condensates described in (**d**). Data are analyzed by one-way ANOVA followed by two-sided Tukey’s multiple comparisons test. Different letters represent significant differences (*P* < 0.05). Data are mean ± SEM (n = 10). **f,** The turbidity of JAZ1 *in vitro*. The absorbance at 600 nm of mCherry-JAZ1 solutions (3 µM) without additives (CK, black line), and with 1 mM of JA-Ile (purple line) or Coronatine (COR, yellow line) were detected. Data are analyzed by one-way ANOVA followed by two-sided Tukey’s multiple comparisons test (n.s: no significance, \*\**P* < 0.01). Data are mean ± SEM (n = 4 biological replicates). **g,** MST assay shows a binding activity of JAZ1 with the Coronatine and JA-Ile but not JA. The binding constant K_D_ values were determined to be 0.17 ± 0.11 μM for Coronatine, and 0.83 ± 0.29 μM for JA-Ile. Green, purple and yellow lines represent JA, JA-Ile and Coronatine (COR). Data are mean ± SD (n = 3 biological replicates).

Coronatine, produced by *Pseudomonas syringae*, has been reported to mimic JA-Ile in activating JA signaling transduction^36^. Notably, we found that both JA-Ile and coronatine facilitates the droplet fusion of mCherry-JAZ1 (Fig. 3d, e). Consistently, the turbidity of mCherry-JAZ1 solutions is gradually increases by both chemicals (Fig. 3f). These results imply that JAZ1 directly perceives JA-Ile and coronatine and undergoes phase separation *in vitro.* This prompts us to propose that the active JA ligand might have a direct interaction with JAZ1. In microscale thermophoresis (MST) assay, the clear binding affinities of mCherry-JAZ1 with coronatine and JA-Ile were observed and the binding constant K_D_ values were 0.17 ± 0.11 μM for coronatine, and 0.83 ± 0.29 μM for JA-Ile (Fig. 3g). When mCherry was used for MST assay, no binding activity between mCherry and coronatine/JA-Ile was found (Extent Data Fig. 10). Taken together, these results indicate that JAZ1 is a direct sensor in JA signaling.

### The ARR residues in Jas domain are required for JAZ1 condensation

Most of the JAZ proteins consist of ZIM (initially named for a zinc-finger protein expressed in the inflorescence meristem) and Jas domains which are essential for their function^1^. The phase separations of the truncated JAZ1 variants in transiently expressed *N. benthamiana* leaves were investigated. We observed that the absence of ZIM domain (JAZ1^δZIM^), the entire C-terminal IDR containing the Jas domain (JAZ1^δIDR^), or the Jas domain (JAZ1^δJas^) results in the elimination of JAZ1 condensation in Zone 2. We further found that deletion of only three residues (ARR) in Jas domain (JAZ1^δARR^) is sufficient for condensation eliminations (Extent Data Fig. 11a, b). The *35S:V-JAZ1^δARR^* transgenic plants under the Col-0 and *jazQ* background were generated. The mRNA and protein levels of JAZ1 in transgenic plants were examined (Extent Data Fig. 12a, b). And we found that JAZ1^δARR^ is unable to form condensates in plant response to either wounding or MeJA treatment (Extent Data Fig. 12c-e). These results indicate that the residues ARR in Jas domain is required for JAZ1condensation.

### JAZ1 switches its binding affinity from MYC2 to COI1 through condensation

The primary transcription factor in JA signaling, MYC2, has been reported to be targeted by JAZ proteins, resulting in the suppression of its transcriptional activity^6^. We examined whether the JAZ1 condensation has an impact on JAZ1-MYC2 interaction. Upon MeJA treatment, V-JAZ1^δARR^ in *N. benthamiana* leaves was more likely to be co-immunoprecipitated with the recombinant MYC2 than V-JAZ1 (Fig. 4a). When the *N. benthamiana* leaves co-expressing with different combinations were treated with MeJA followed by a dual-luciferase reporting system assay, we observed that compared to V-JAZ1, V-JAZ1^δARR^ exhibits a stronger inhibition on MYC2-mediated transcriptional activation (Fig. 4b). This reveals that JAZ1 shows higher binding affinity with MYC2 in dispersed than condensed phase.

**Fig. 4:**
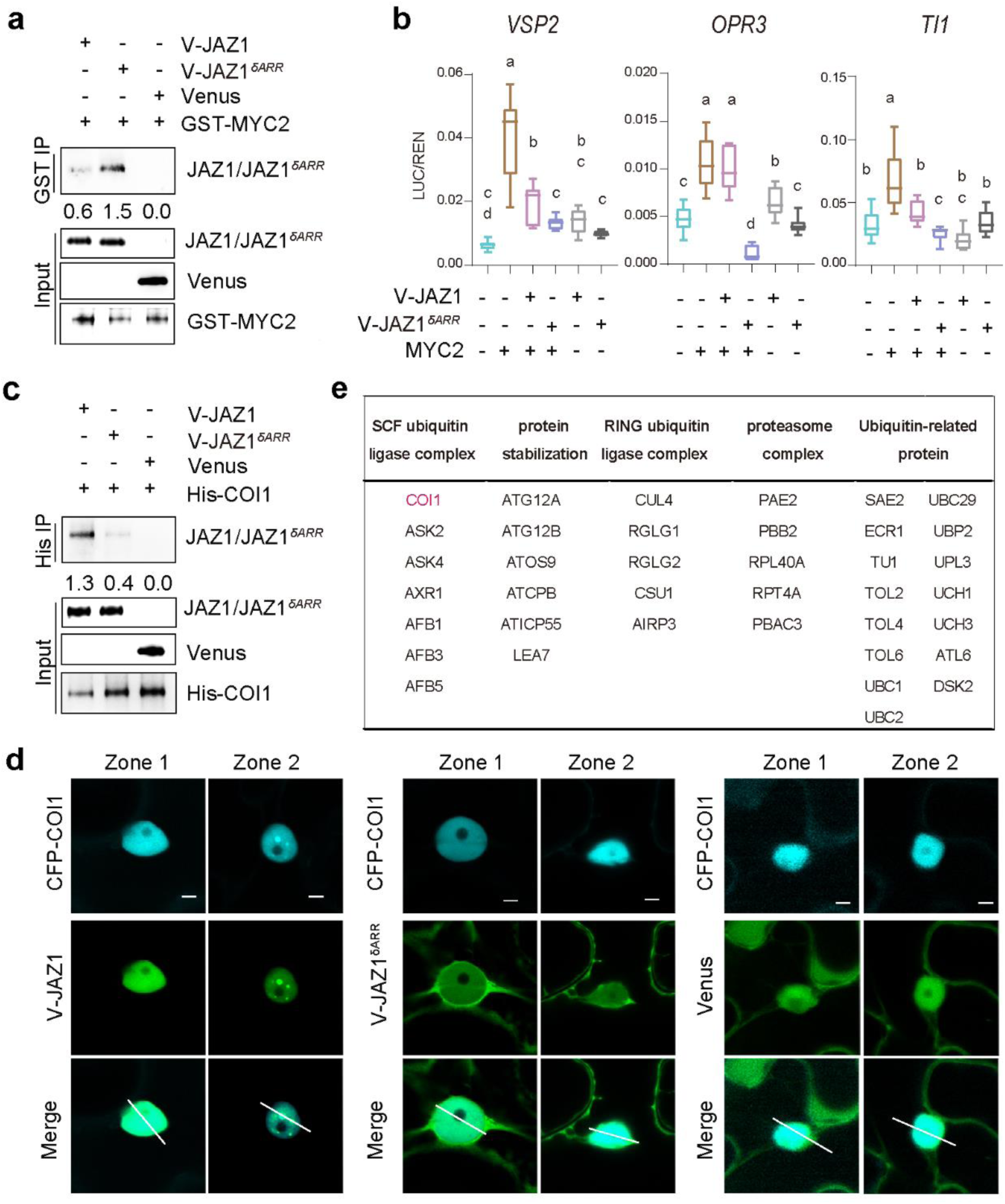
Phase separation regulates JAZ1 affinity with MYC2 and COI1 in opposite way. **a,** MYC2 is inclined to interact with JAZ1 in dispersed phase. The recombinant GST-MYC2 protein is incubated with the total protein extractions of *N. benthamiana* leaves transiently expressed V-JAZ1, V-JAZ1^δARR^ and Venus, respectively. Anti-GST antibody is to detect GST-MYC2 in input. Anti-Venus antibody is used to detect V-JAZ1, V-JAZ1^δARR^ and Venus. **b,** The transcriptional activity of MYC2 is more suppressed by V-JAZ1^δARR^ than those by V-JAZ1. The promoters of MYC2-targeted genes (*VSP2*, *OPR3* and *TI1*) with predicted G-BOX in *Arabidopsis* are fused to the LUC reporter and transiently expression in *N. benthamiana* leaves. The luciferase (LUC) activities were quantified. Data are analyzed by one-way ANOVA followed by two-sided Tukey’s multiple comparisons test. Different letters represent significant differences (*P* < 0.05). The top and bottom lines of the box plot represent the 25th and 75th percentiles, the center line is the median, and the bar shows the range of the highest and lowest data points (n = 8 biological replicates). **c,** Condensed JAZ1 shows higher affinity with COI1 than those in dispersed phase. The recombinant His-COI1 proteins are incubated with the total protein extractions of *N. benthamiana* leaves transiently expressing V-JAZ1, V-JAZ1^δARR^ and Venus, respectively. Pull-down fractions are examined by immunoblot analysis using anti-Venus antibodies to detect V-JAZ1, V-JAZ1^δARR^ and Venus. Anti-His antibody is to detect COI1 in input. **d,** COI1 is incorporated into JAZ1 condensates in plant response to wounding. V-JAZ1, V-JAZ1^δARR^ and Venus were transiently co-expressed with CFP-COI1 in *N. benthamiana* leaves respectively. CFP-COI1 proteins form condensates at Zone 2 in the presence of V-JAZ1 rather than V-JAZ1^δARR^. Scale bars, 5 μm. **e,** Proteins identified by IP-MS in the JAZ1-enriched condensates.

On the contrary, V-JAZ1 was more likely to be co-immunoprecipitated with recombinant COI1 than V-JAZ1^δARR^ from total proteins of *N. benthamiana* leaves which were pretreated with MeJA (Fig. 4c). When CFP-or Venus-fused COI1 were expressed in the *N. benthamiana* leaves, no condensate was observed in both Zone 1 and 2. While co-expressed with V-JAZ1, CFP-COI1 is recruited into JAZ1 condensates in Zone 2 (Fig. 4d and Extent Data Fig. 13a, b). To have a deep insight into the protein components of JAZ1 condensates, we performed the proteomics analysis using affinity-purified JAZ1 condensates. ASK2, ASK4, CUL4, and 36 other proteins related to proteasome degradation were co-purified with JAZ1 condensates (Fig. 4e).

### Invalidating JAZ1 condensation dulls JA signaling transduction

Through single-cell time-lapse imaging, we observed rapid condensation of V-JAZ1 in *35S:V-JAZ1* plants (line 98) upon MeJA treatments, which was initially evident at 5 minutes, became more pronounced at 10-15 minutes, and fade away within 20 minutes. On the other hand, although the fluorescent intensity reduced to some extent in *35S:V-JAZ1^δARR^* plants (line 32) upon MeJA addition, V-JAZ1^δARR^ was unable to form condensates (Fig. 5a and Supplementary Video 9, 10). The quantification of total fluorescent intensity reveals a more rapid decrease of V-JAZ1 compared to V-JAZ1^δARR^ (Fig. 5b). Immune blot assay also showed that the accumulation of V-JAZ1 in *35S:V-JAZ1* plants and V-JAZ1^δARR^ in *35S:V-JAZ1^δARR^* plants were similar in the absence of MeJA, however, upon MeJA treatment, the decrease of V-JAZ1 was more pronounced compared to that of V-JAZ1^δARR^ (Extent Data Fig. 14a, b). These results suggest that V-JAZ1 exhibits a higher sensitivity to degradation in plant response to MeJA compared to V-JAZ1^δARR^.

**Fig. 5:**
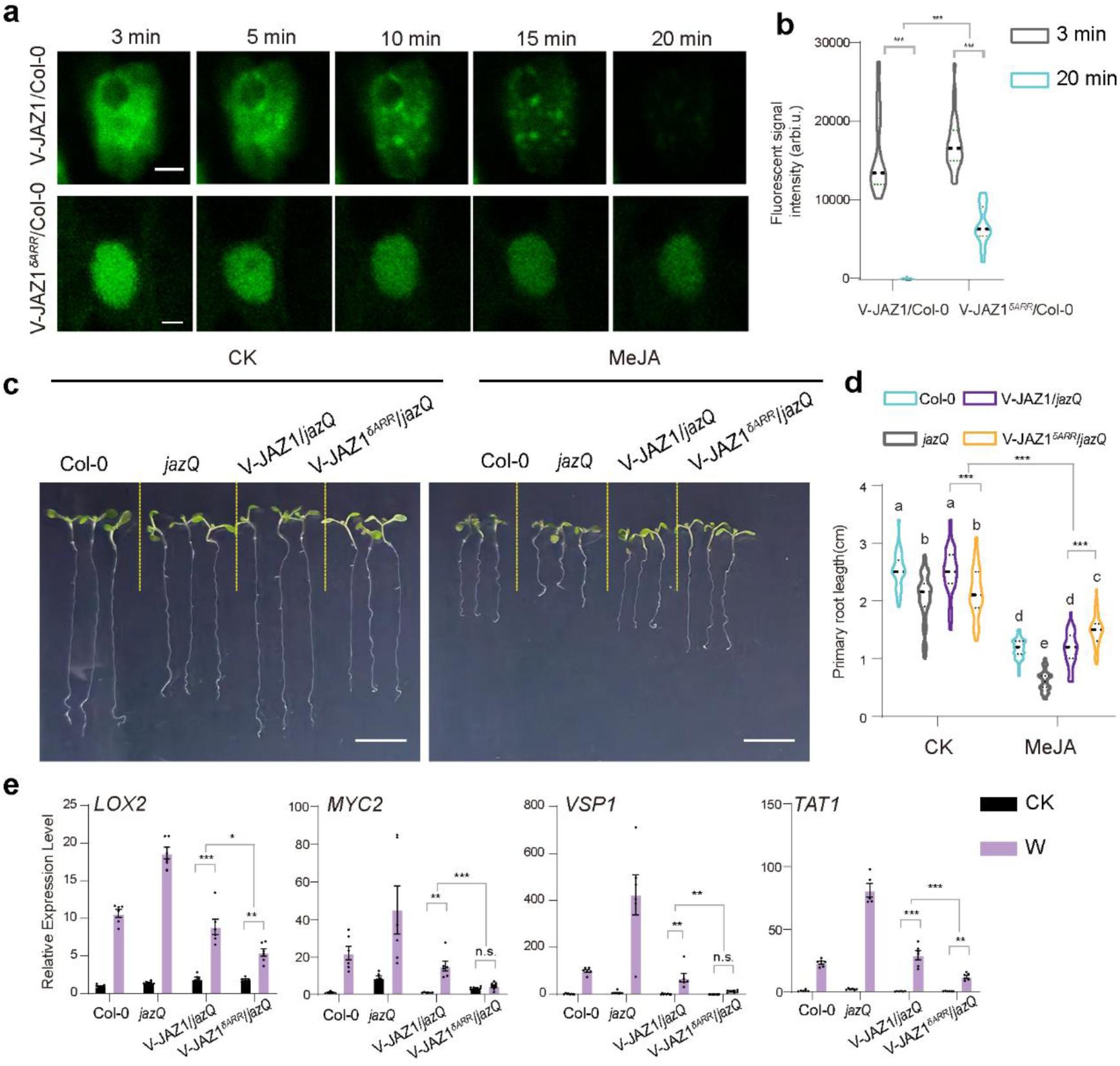
The ability of JAZ1 to form condensates enables a quick JA signaling transduction in plant. **a-b,** Condensation facilitates the degradation of JAZ1. **a,** Time-lapse imaging of V-JAZ1 and V-JAZ1^δARR^ post 50 μM MeJA treatment in *35S:V-JAZ* line 98 (V-JAZ1/Col-0) and *35S:V-JAZ1^δARR^* line 32 (V-JAZ1^δARR^/Col-0) transgenic plants. Images are started immediately after MeJA addition (3 min) for a duration of 20 min. Images from selected time points were presented. Scale bars, 5 μm. **b,** Quantification of JAZ1 fluorescence intensity (arbitrary units, arbi.u.) 3 min and 20 min post MeJA treatment as described in (**a**). 30 random nuclear from 5 plants were selected for calculation. Data are analyzed by two-way ANOVA followed by multiple comparisons with two-sided Fisher’s LSD test (\*\*\**P* < 0.001). Different letters represent significant differences (*P* < 0.05). Data are mean ± SEM (n = 30). Brown and blue represent 3 min and 20 min respectively. **c-d,** *35S:V-JAZ1 jazQ* exhibits a JA response comparable to the wild type, whereas *35S:V-JAZ1*^δARR^ *jazQ* is insensitive to JA. **c,** Inhibition of JA on the root growth. The seeds of *jazQ*, *35S:V-JAZ1 jazQ* line 1 (V-JAZ1/*jazQ*) and *35S:V-JAZ1*^δARR^*jazQ* line 15 (V-JAZ1^δARR^/*jazQ*) were germinated on the 1/2 MS supplemented with 5 μM MeJA. The 7-day-old seedlings were analyzed. Scale bars, 1 cm. **d,** Quantification of the root length as described in (**c**). Data are mean ± SEM (n = 85 biological replicates) and analyzed by two-way ANOVA followed by multiple comparisons with two-sided Fisher’s LSD test (n.s.: no significance, \*\*\**P* < 0.001). **e,** qRT-PCR analysis of gene expressions. 17-day-old plants were wounded (W), and leaves were collected 2 h after treatment. Leaves from the unwounded plants were used as control (CK). Data are analyzed by two-way ANOVA followed by multiple comparisons with two-sided Fisher’s LSD test (n.s.: no significance, \**P* < 0.05, \*\**P* < 0.01, \*\*\**P* < 0.001). Data are mean ± SEM (n = 6 biological replicates).

The degradation of JAZ proteins determines the output of JA signaling. When grown on the normal plant culture medium, the root of *jazQ* seedlings exhibits a stunted growth compared to the wild-type Col-0. And this phenotype is restored in *35S:V-JAZ1 jazQ* whereas unchanged in *35S:V-JAZ1^δARR^ jazQ*. Upon MeJA treatment, root growth was inhibited and the inhibition is similar between Col-0 and *35S:V-JAZ1 jazQ* while more severe in *jazQ.* Notably, the slightest inhibition on root growth was observed from the *35S: V-JAZ1^δARR^ jazQ* plant suggesting it is insensitive to MeJA treatment (Fig. 5c, d). Wounding quickly elicits a large amount of JAs in plants and induces defense gene expressions. We found *jazQ* exhibits the highest inductions of *LOX2*, *MYC2*, *VSP1* and *TAT1* expression, and the inductions are comparable between wild type and *35S:V-JAZ1 jazQ* plants suggesting that the hypersensitive response to wounding in *jazQ* is restored by the expression of *V-JAZ1.* However, the inductions are significantly compromised in *35S:V-JAZ1^δARR^ jazQ* plants compared to wild-type and *35S:V-JAZ1 jazQ* plants (Fig. 5e). These results indicated that phase separation of JAZ1 is essential for its proper functioning in signaling transduction in wounding stress.

## Discussion

JAZ proteins play pivotal roles as central molecular hubs, integrating a wide range of signals and orchestrating responses that profoundly influence plant development, defense against biotic and abiotic stresses^37^. We found that JAZ proteins undergo condensations via LLPS upon wounding (Fig. 1). In this process, the active JA ligand is required for wounding-triggered JAZ1 condensates (Fig. 2, 3). The interaction between MYC2 and JAZ1 is more likely to occur in the dispersed phase rather than the condensed phase, while COI1 is more inclined to interact with condensed JAZ1 in the presence of JA-Ile (Fig. 4). Without wounding, the contents of JAs are at low level. And JAZ proteins are dispersed in nucleus, it interacts with MYC2 and blocks the signaling output. Upon wounding, JAs are quickly elicited. JAZ1 directly binds with JA-Ile, causing a transition from a dispersed to condensed phase. JAZ1 condensates recruit the 26S proteasome related proteins for efficient degradation. Thereby, the rapid switch of JA signaling from a stationary to an active state is achieved by the phase transitions of JAZs (Fig. 6). Furthermore, the *35S:V-JAZ1^δARR^ jazQ* plant exhibits a stunted root growth like *jazQ* in basal condition (Fig. 5) suggesting that the function of JAZ condensation is beyond wounding and JA perceptions.

**Fig. 6:**
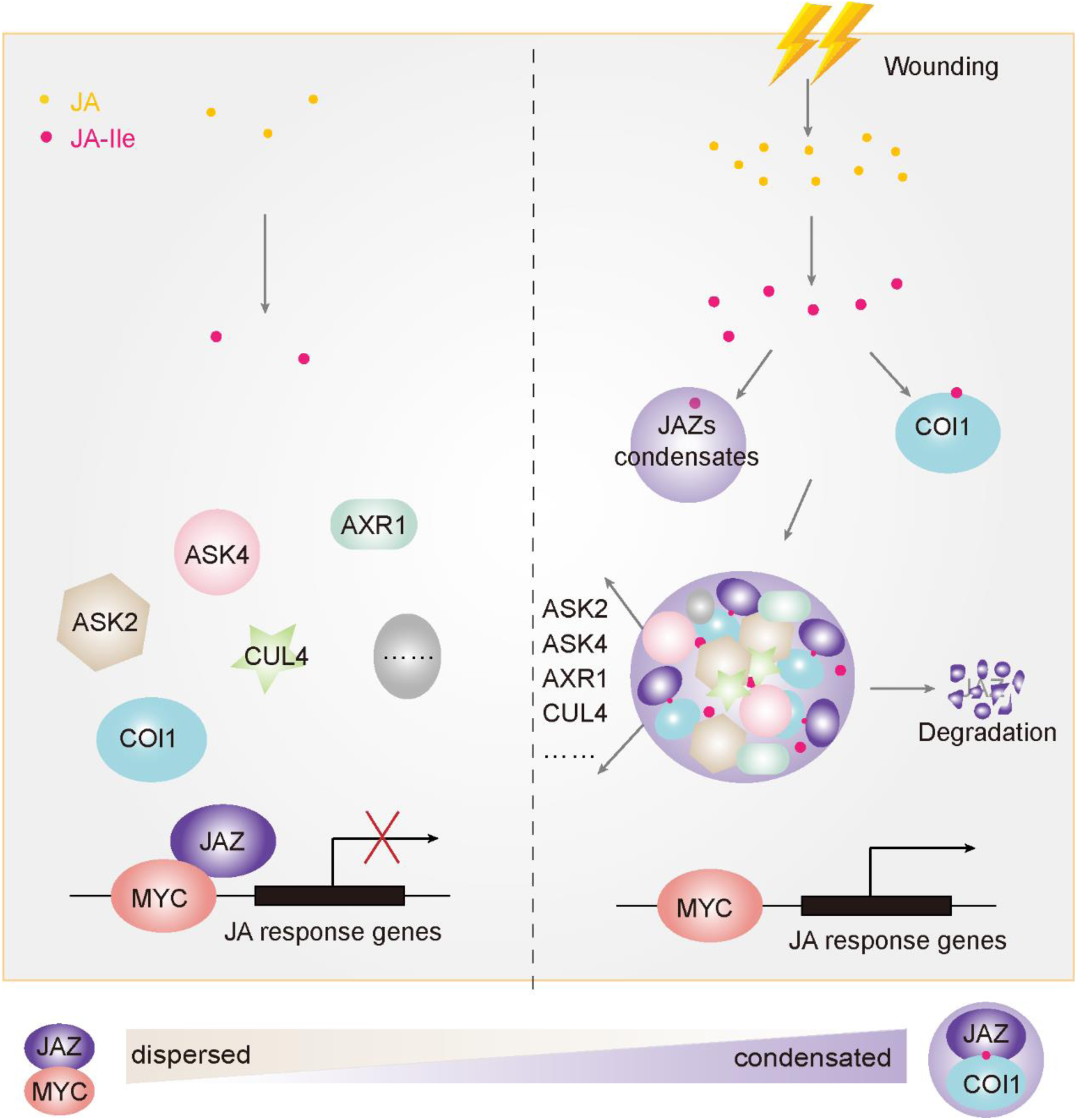
Diagram of JAZs regulating JA signaling output by switching from dispersed to condensed phase. In basal condition, JAZ proteins are in dispersed phase and bind with MYC2 to inhibit downstream gene expression. JAZs independently perceive JA-Ile which is elicited by wounding and undergo condensation. The condensed JAZs reduced their binding affinity with MYC2 and tend to recruit COI1 and other proteasome related proteins to achieve quick JAZ degradation thereby prompting the signaling transduction.

The IDR plays crucial roles in diverse biological processes and pathways, and it is also required for condensate formation under stress conditions^38^. Besides multiple JAZ proteins of Arabidopsis contain IDR (Extent Data Fig. 1b), JAZ proteins in *Marchantia polymorpha* (liverwort), *Huperzia selago* (lycophyte), *Oryza sativa* (monocot) plants reveal a common feature of the presence of IDRs (Extent Data Fig. 2). This suggests that IDR within JAZ proteins have been evolutionarily conserved. Interestingly, the mutation of three residues (ARR) in Jas domain which imbedded in IDR abolishes JAZ1 condensations upon JA perception. Together with the observation of the clear direct interaction between JAZ1 and JA-Ile (Fig. 3), it can be inferred that ARR in jas is essential for establishing a strong binding affinity with the active ligand. Evidences have shown that in some cases, IDR alone may not be sufficient for condensation, but combined with specific interactions to selectively drive biomolecular condensation^39^. Here, we found that JAZ1^δZIM^ completely lost the ability to form condensates (Extended Data Fig. 11). ZIM domain has been reported to mediate JAZ proteins homo and hetero dimerization and also involved in the recruitment of TPL and TPR proteins indirectly through the adaptor, NINJA^1^. Our study reveals that ZIM domain coordinates with IDR to fulfill a crucial role in protein phase separation.

The initial step in signal transduction involves the specific recognition of the active ligand by its receptor. JA-Ile has been known to be perceived by COI1-JAZ coreceptors. And the interaction between COI1 and JA-Ile exhibits enhanced stability in the presence of JAZ proteins^11^. Besides, JAZ proteins also contributes to determine the specificity of the ligands^12^. However, only COI1 has been verified for its direct interaction with active ligand^13^. It is widely accepted that COI1 primarily senses active JA ligand with the assistance of JAZs. In this study, we have shown that JAZ1 exhibit a direct binding affinity with JA-Ile and coronatine in MST assay (Fig. 3). JAZ1 switches from dispersed to condensed phase in the presence of active ligands while the mutated JAZ1 (JAZ1^ARR^) is deficient in phase transition (Fig. 5). Together, we initially provide evidence that JAZ1 functions as a direct independent sensor in JA signaling, regardless of COI1. The perception of JA-Ile by COI1 triggers its interaction with JAZ proteins leading to the subsequent degradation of the latter through ubiquitin-dependent 26S proteasome for signaling initiation. This function positions COI1 as a “starter” that initiates the cascade of signaling events. On the other hand, JAZs can independently sense JA-Ile via phase separation, the condensed JAZ1 decreases its interaction with MYC2 and recruits 26S proteasome related components to achieve rapid degradation. In this way, the condensation of JAZ can be regarded as an “accelerating process” of the signaling output. This offers a possibility that differential responses of plant to distinct stimuli can be achieved by modulating JAZ condensations. Together, COI1 is responsible for initiating the signaling process, whereas JAZ controls the rate of signaling transduction. The dual sensor system enables a more sophisticated regulation and an expanded repertoire of responses to intricate stimuli, ultimately facilitating effective defense.

## Methods Details

### Plant materials, culture and treatment

All the *Arabidopsis thaliana* plants used in this study were of the Columbia, Col-0 ecotypes. The mutants, *jazQ*^7^, *aos*^33^ and *coi1-2*^35^ were described previously. Transgenic plants were generated in the laboratory, with wild type (Col-0) background (*35S:Venus-JAZ1*, *35S:Venus-JAZ1^δARR^*, *35S:Venus*), *jazQ* background (*35S:Venus-JAZ1 jazQ*, *35S:Venus-JAZ1^δARR^ jazQ*), *aos* background (*35S:Venus-JAZ1 aos*) and *coi1-2* background (*35S:Venus-JAZ1 coi1-2*).

All the plants were grown under long photoperiod conditions (16 h light/8 h dark, 22°C). The surface-sterilized seeds of *Arabidopsis* were kept at 4°C in the dark for 2 days before being used for germination on half-strength Murashige and Skoog (1/2 MS) medium (pH 5.8) containing 0.75% agar and 1% sucrose. Seedlings were transferred on soil, 7 days post germination.

To examine the gene expressions in wounding treatment, about 1/3 areas of the leaves from 17-day-old plants were punched and the unwounded leaves were used as control. Two hours later, the leaves were collected. And the total RNA extractions were used for qRT-PCR analysis. To observe the wounding induced condensation, leaves or roots of 6-day-old transgenic plants were wounded by nippers. The leaf pavement cells and root elongation region cells were immediately collected for confocal microscopy.

For MeJA experiments, MeJA (Sigma-Aldrich, 392707) was dissolved in ethanol and diluted in ddH_2_0 to a final concentration of 50 μM. As the mock treatment, equal volumes of ethanol were used. The leaves were sprayed with water solutions of MeJA and ethanol, respectively. The leaves were collected immediately for confocal microscopy.

### Plant transformation

The coding sequences (CDSs) of *COI1* (*AT2G39940*), *JAZ1* (*AT1G19180*) and truncated *JAZ1* were amplified by PCR from Col-0 cDNA. The truncated *JAZ1* variants in extent data figure 10a were synthesized by Tsingke (Beijing, China). COI1 with CFP fused in N-terminal, JAZ1 and its truncated variants with Venus fused in N-terminal as well as Venus and CFP were expressed under cauliflower mosaic virus 35S promoter. Vector constructions were performed by homologous recombination with ClonExpress Ultra One Step Cloning Kit (Vazyme, C115-01). All the constructed vectors were transformed into *Agrobacterium tumefaciens* strain GV3101 (WEIDI, AC1001) by the freeze–thaw method. The primers are provided in Supplementary Table S1.

For transient expression in the leaves of *N. benthamiana*, the *A. tumefaciens* cells carrying the desired construct were cultured overnight at 28°C in Luria–Bertani broth supplied with kanamycin (50 mg/L), gentamicin (50 mg/L) and rifampicin (25 mg/L), then were transferred to the new Luria– Bertani broth at a ratio of 1:500 and cultured overnight. The *A. tumefaciens* suspension cultures were centrifuged at 3,000 *g* for 5 min and the precipitates were re-suspended in infiltration buffer (10 mM MgCl2, 10 mM MES, 150 μM Acetosyringone, pH = 5.7) with an OD600 around 1.0. After incubated at room temperature for 3 h, the cell resuspensions were infiltrated into leaves of four-week-old *N. benthamiana* with 1 ml syringes. 2-3 days later, leaves were harvested for pull-down and microscopy experiments.

The flower dipping method was used for generating the transgenic *Arabidopsis*^40^. The inflorescence of *Arabidopsis* plants was soaked in a suspension of *A. tumefaciens* cells carrying the desired construct. Seeds of the T0 generation were screened on 1/2 MS medium containing 50 μg/ml Hygromycin B. 8 days post germination, the positive seedlings were transferred to soil. The expression of transgene was further examined by qRT-PCR and immune blot.

### Prokaryotic Expression and Purification

For expression of His-fused proteins (His-mCherry-JAZ1 and His-COI1), the corresponding sequences were inserted into the pET-Duet vector. For expression of GST-fused MYC2, the CDS of *MYC2* (*AT1G32640*) was inserted into pGEX-4T-1 vector with a GST N-terminal fusion. The primers are listed in Supplementary Table 1.

*Escherichia coli* strain BL21 (WEIDI, C504) was used to express recombinant protein. Cells carrying the desired construct were cultured at 37°C until they reached an optical density of 1.0 at 600 nm. And then, β-D-1-thiogalactopyranoside (IPTG) was added to a final concentration of 0.25 mM to induce expression. After incubating at 16 °C overnight, cells were collected for protein purification. Ni affinity column (Ni-NTA resin, Qiagen) and Glutathione Sepharose 4B resin (GEHealthcare) were used for the purification of His-fused proteins and the GST fusion proteins respectively. The eluted proteins were concentrated and desalted using an Amicon Ultra-15 Centrifugal Filter Unit (EMD Millipore, UFC901096-1) with 50 mM Tris– HCl (pH = 8.0). All proteins were quantified and then aliquoted for storage at −80°C.

### *In vitro* droplet assay

The purified His-mCherry-JAZ1 and His-mCherry were diluted to the desired concentrations using 50 mM Tris-HCl (pH=8.0). To induce phase separation, crowding agent PEG8,000 (Beyotime, ST484) was added to a final concentration of 10% (w/v) and thoroughly mixed followed by confocal microscopy observation. To examine the impacts of JA-Ile (GLPBio, GC19497-1) and Coronatine (Sigma, C8115) on JAZ1 condensation, these two chemicals were respectively added to the JAZ1 solutions to a final concentration of 1 mM before the confocal microscopy observation.

### Turbidity measurement

The recombinant mCherry-JAZ1 protein was prepared in 50 mM Tris-HCl (PH=8.0) containing 10% PEG (w/v) to a final concentration of 3 μM. JA-Ile and Coronatine stocked in 50 mM Tris-HCl was added to the mCherry-JAZ1 solution to a final concentration of 1 mM. The mCherry-JAZ1 solution added with equal volume of 50 mM Tris-HCl was used as control (CK). Turbidity was measured at the optical absorption of 600 nm every 2 min for 4 h by a multisacan GO (Thermo Fisher) using a ANS/SBS standard plate 96-well plates.

### Microscale Thermophoresis Assay (MST)

mCherry-JAZ1 was NHS labeled and kept in 50 mM Tris-HCl (PH=8.0). The substrates Coronatine, JA-Ile and jasmonate acid were stocked in PBST (PBS containing 0.02% TWEEN). NHS-labeled mCherry-JAZ1 of 20 nM were mixed with equal volume of substrate at different concentrations from 100 μM to 0.00305 μM. After brief incubation, the samples were loaded into MST-standard glass capillaries and measured by the Monolith NT.115 (MicroScale) at 24℃. Data analyses were performed through the Nanotemper Analysis and MO. Affinity.

### Confocal microscopy

All images and movies were taken by confocal systems, Leica SP8, Leica STELLARIS5 or Nicon AX equipped with different immersive objectives (Leica SP8 with ×20 and ×63 water-immersion objective, Leica STELLARIS5 with ×20 water-immersion objective and ×63 oil-immersion objective, Nicon AX with ×100 oil-immersion objective). For time-lapse microscopy observation, images were acquired every 1 second for 2 min (*in vitro*) or every 5 s for 2 min (*in vivo*).

The excitation/emission wave lengths CFP, Venus and mCherry signals were 405/450-500, 488/490-579 and 561/610-650 nm, respectively. And all images were acquired using Leica or Nicon software. The fluorescence signal intensity was calculated using ImageJ software.

### Quantification of JAZ1 condensates

The circularity and diameter quantifications of JAZ1 condensates were analyzed by FIJI/ImageJ as previously described^41^. In brief, the images use the “Process” tool to set up the subtract background. Then the “Image” tool was used to adjust the threshold. Each individual condensate was analyzed by “Analyze particles” to determine the circularity and diameter.

### Fluorescence recovery after photobleaching (FRAP) assay

JAZ1 condensation was observed under a Leica TSC SP8 confocal microscopy. The selected JAZ1 condensate was first recorded with 3 iterations, and then was bleached with 5 iterations using a laser intensity of 100% at 561 nm for mCherry-JAZ1 condensate in vitro and at 488 nm for Venus-JAZ1 condensate in vivo. After bleaching, the fluorescence recovery of *in vitro* and *vivo* condensates ware recorded at intervals of every 1 s and every 3 s respectively, for 1 min.

Fiji/ImageJ was used to analyze the fluorescence intensity of bleached region, reference region and background region. Recovery curves were drawn using the web-based tool easyFRAP (http://easyfrap.vmnet.upatras.gr).

### Pull-down assay

To detect the interactions of JAZ1 and JAZ1^δARR^ with His-COI1 or GST-MYC2, the *N. benthamiana* leaves transiently expressing Venus and Venus fused JAZ1/JAZ1^δARR^ were treated with 50 μM MeJA and the total leaf proteins were extracted from the samples 30 min post treatment. His-COI1 (20 μg) was incubated with the protein extractions in the presence of 10 μL Ni while GST-MYC2 (20 μg) was incubated with the protein extractions in the presence of 10 μL GST affinity beads, at 4°C with rotation for overnight.

### Immune blot analysis

Proteins were separated by 10% or 12.5% SDS-PAGE and transferred onto first antibodies. Anti-Venus antibody (ABclonal, AE012, 1:2,000) was used to detect Venus, Venus-JAZ1 and Venus-JAZ1^δARR^. Anti-His antibody (ABclonal, AE003, 1:2,000) was used to detect COI1 in input. Anti-GST (ABclonal, AE001, 1:2,000) antibody was used to detect GST-MYC2 in input. After incubation with first antibodies, the blots were incubated with horseradish-peroxidase-conjugated anti-mouse secondary antibody (ABclonal, AS003, 1:2,000) or anti-rabbit secondary antibody (ABclonal, AS014, 1:2,000). After the addition of ECL prime western blot detection reagent (Epizyme, SQ201), chemiluminescence was detected using Image Lab (Bio-Rad).

### Dual-luciferase assay

The 500-2000 bp upstream sequence of the start codon of *VSP2* (*AT5G24770*), *OPR3* (*AT2G06050*) and *TI1* (*AT2G43510*) genes was used as the promoter and inserted into the pGreen-LUC vector to drive firefly LUC reporter gene expression. Renilla (REN) driven by 35S promoter was used for normalization. Agrobacterium strains harboring *35S:MYC2-Flag* were used as activators, and Venus-JAZ, Venus-JAZ1^δARR^ and Venus (negative control) were added as repressors. Agrobacterium strains with the indicated promoters, activators, repressors were mixed and injected into tobacco leaves. A dual-luciferase reporter assay system (Promega, E1910) was used for protein extraction and enzyme activities examination. Fluorescence values of LUC and REN were detected using a Varioskan Flash (Thermo). The ratio of LUC : REN was presented as promoter activity. Eight biological replicates were measured for each data point.

### JAZ1 condensates isolation

Condensates were isolated from the 8-day-old seedlings of *35S:Venus-JAZ1* and *35S:Venus-JAZ1^δARR^* (negative control) using the previously described method^42^. The seedlings were treated with 50 μM MeJA and 10 μM MG132 simultaneously and harvested 2 h post treatment. 2 g of seedlings were ground into fine powder in liquid nitrogen. The ground powder was then added to 2 mL of ice-cold lysis buffer (50 mM Tris–HCl pH 7.4, 100 mM potassium acetate, 2 mM magnesium acetate, 0.5% NP-40, 0.5 mM DTT, 1 mM NaF and 10 μM MG132) and incubated on ice for 10 min. The lysate was centrifuged at 4,000 g for 10 min at 4 °C. The pellet was resuspended in 1 ml of lysis buffer, vortexed, and centrifuged at 18,000 g for 10 min at 4 °C. Repeat this step once. The pellet was then resuspended in 1 ml of lysis buffer and centrifuged at 850 g for 10 min at 4 °C. The supernatant was checked for the presence of JAZ1 condensates by a Leica STELLARIS5 confocal microscope.

### IP-MS analysis

The extracted condensate fraction was used for IP-MS with Flag-Catch-Mag Beads (Engibody, AT1770). The JAZ1 condensate-enriched fraction were incubated with 25 μl of Flag-Catch-Mag beads for overnight at 4 °C. Samples on beads were reduced with 10 mM dithiothreitol for 30 min at 55 °C followed by alkylation with 30 mM iodoacetamide in dark for 15 min at room temperature. Then 1 mg trypsin (Promega, V5111) was added for overnight digestion at 37 °C. After desaltination, the samples were dried in SpeedVac. Peptides were redissolved in 0.1% FA and 1 mg sample was loaded and analyzed with timsTOF Pro 2 (Bruker). Mass spectrometry analyses were performed at the Proteomics Facility of CAS center for excellence in molecular plant sciences. The total of proteins were detected in the IP fractions of *35S:Venus-JAZ1* samples in Supplementary Table 2.

### Quantitative analysis of root length

MeJA was dissolved in 1% (v/v) ethanol as a 50 mM stock solution. Arabidopsis seeds were germinated and grown for 7 d on 1/2MS medium supplemented with 5 μM MeJA. In the mock treatment, an equal volume of ethanol was added to the medium.

### Gene expression analyses

Total RNA was extracted from the second pair of true leaves of a 17-day-old plant using Trizol reagent (Invitrogen, 15596018). The cDNA was reverse transcribed from 1.5 ug total RNA with oligo (dT) primers using TransScript One-Step gDNA Removal and cDNA Synthesis SuperMix kit (TransGen, AT311-03). The real-time-PCR was performed with SYBR green PCR master mix (TaKaRa, QPK-201C) on a real-time PCR system (CFX thermocycler; Bio-Rad, Hercules, CA). For data analyses, we used the delta CT (critical threshold) method and the mRNA level of S18 (At4g09800) was used as an internal reference. The primer sequences used are listed in Supplementary Table 1.

### Statistical analysis

For parametric statistics, a normal distribution was assumed if the P value of the Shapiro-Wilk test was larger than 0.05. Two-sided Student’s t-tests, one-way or two-way analysis of variance ^43^ tests were used for two and multiple comparisons, respectively. Statistical tests were performed in GraphPad Prism 8 with P > 0.05 considered not significant. Unless specifically stated, sample size n means that the biological replications and experiments have been performed at least three times (one time for LC-MS).

## Acknowledgements

We thank Prof. Xiao-Ya Chen and Peng Zhang (CEPMS/SIPPE, CAS) for helpful discussions and suggestions, Li Liu (CEPMS/SIPPE, CAS) for MST assay of small molecule-protein interaction, Shanshan Wang, Lianyan Jing and Zhihui Wang (CEPMS/SIPPE, CAS) for their assistance in identification of condensate proteins. This research was supported by National Key R&D Program of China (2023ZD04073), National Natural Sciences of China Grants (32072430 and 32000221), Science and Technology Commission of Shanghai Municipality (22ZR1469300 and 21ZR1471000).

## Author contributions

Z.W.Y. and Y.B.M. designed the research. Z.W.Y. performed the main experiments with the assistances from J.L., H.C.C., J.X.L., M.Y.W and W.J.C. Data were analyzed by Z.W.Y., J.W.W. and Y.B.M. The manuscript was written by Z.W.Y. and Y.B.M. with input from all authors.

## References

1 Howe, G. A., Major, I. T. & Koo, A. J. Modularity in Jasmonate Signaling for Multistress Resilience. Annu Rev Plant Biol 69, 387–415 (2018). 10.1146/annurev-arplant-042817-040047

2 Erb, M. & Reymond, P. Molecular Interactions Between Plants and Insect Herbivores. Annu Rev Plant Biol 70, 527–557 (2019). 10.1146/annurev-arplant-050718-095910

3 Chen, X., Wang, D.-D., Fang, X., Chen, X.-Y. & Mao, Y.-B. Plant Specialized Metabolism Regulated by Jasmonate Signaling. Plant and Cell Physiology 60, 2638–2647 (2019). 10.1093/pcp/pcz161

4 Wasternack, C. & Song, S. Jasmonates: biosynthesis, metabolism, and signaling by proteins activating and repressing transcription. J Exp Bot 68, 1303–1321 (2017). 10.1093/jxb/erw443

5 Browse, J. Jasmonate Passes Muster: A Receptor and Targets for the Defense Hormone. Annual Review of Plant Biology 60, 183–205 (2009). 10.1146/annurev.arplant.043008.092007

6 Zhang, F. et al. Structural basis of JAZ repression of MYC transcription factors in jasmonate signalling. Nature 525, 269–273 (2015). 10.1038/nature14661

7 Campos, M. L. et al. Rewiring of jasmonate and phytochrome B signalling uncouples plant growth-defense tradeoffs. Nature Communications 7 (2016). 10.1038/ncomms12570

8 Yan, C. et al. Injury Activates Ca(2+)/Calmodulin-Dependent Phosphorylation of JAV1-JAZ8-WRKY51 Complex for Jasmonate Biosynthesis. Mol Cell 70, 136–149 e137 (2018).10.1016/j.molcel.2018.03.013

9 Xie DX, F. B., James S, Nieto-Rostro M, Turner JG. May COI1: an Arabidopsis gene required for jasmonate-regulated defense and fertility. Science. (1998).

10 A. Chini1*, S. F., G. Fernández1*, B. Adie1, J. M. Chico1, O. Lorenzo1{, G. Garcìa-Casado2, I. López-Vidriero2, F. M. Lozano3, M. R. Ponce3, J. L. Micol3 & R. Solano1,2. The JAZ family of repressors is the missing link in jasmonate signalling. nature (2007). 10.1038/nature06006

11 Laura B. Sheard, X. T., Haibin Mao, John Withers, Gili Ben-Nissan, Thomas R. Hinds, Yuichi Kobayashi, Fong-Fu Hsu, Michal Sharon, John Browse, Sheng Yang He, Josep Rizo, Gregg A. Howe & Ning Zheng. Jasmonate perception by inositol-phosphate-potentiated COI1– JAZ co-receptor. (2010). 10.1038/nature09430

12 Isabel Montea, J. C., Angel M. Zamarreñoc, Gemma Fernández-Barberoa, José M. García-Minac, and Roberto Solanoa,. JAZ is essential for ligand specificity of the COI1/JAZ co-receptor. Proceedings of the National Academy of Sciences (2022). 10.1073/pnas

13 Hu, S., Yu, K., Yan, J., Shan, X. & Xie, D. Jasmonate perception: Ligand–receptor interaction, regulation, and evolution. Molecular Plant 16, 23–42 (2023). 10.1016/j.molp.2022.08.011

14 Yan, J. et al. Dynamic Perception of Jasmonates by the F-Box Protein COI1. Molecular Plant 11, 1237–1247 (2018). 10.1016/j.molp.2018.07.007

15 Brangwynne, C. P. et al. Germline P granules are liquid droplets that localize by controlled dissolution/condensation. Science 324, 1729–1732 (2009). 10.1126/science.1172046

16 Banani, S. F., Lee, H. O., Hyman, A. A. & Rosen, M. K. Biomolecular condensates: organizers of cellular biochemistry. Nat Rev Mol Cell Biol 18, 285–298 (2017). 10.1038/nrm.2017.7

17 Liu, X., Zhu, J.-K. & Zhao, C. Liquid-liquid phase separation as a major mechanism of plant abiotic stress sensing and responses. Stress Biology 3 (2023). 10.1007/s44154-023-00141-x

18 Zhu, S. et al. Liquid-liquid phase separation of RBGD2/4 is required for heat stress resistance in Arabidopsis. Dev Cell 57, 583–597 e586 (2022). 10.1016/j.devcel.2022.02.005

19 Chen, D. et al. Integration of light and temperature sensing by liquid-liquid phase separation of phytochrome B. Molecular Cell (2022). 10.1016/j.molcel.2022.05.026

20 Zhu, P., Lister, C. & Dean, C. Cold-induced Arabidopsis FRIGIDA nuclear condensates for FLC repression. Nature 599, 657–661 (2021). 10.1038/s41586-021-04062-5

21 Zavaliev, R., Mohan, R., Chen, T. & Dong, X. Formation of NPR1 Condensates Promotes Cell Survival during the Plant Immune Response. Cell 182, 1093–1108 e1018 (2020). 10.1016/j.cell.2020.07.016

22 Wang, J. et al. Phase separation of nuclear pore complex facilitates selective nuclear transport to regulate plant defense against pathogen and pest invasion. Mol Plant (2023). 10.1016/j.molp.2023.04.008

23 Wright, P. E. & Dyson, H. J. Intrinsically disordered proteins in cellular signalling and regulation. Nature Reviews Molecular Cell Biology 16, 18–29 (2014). 10.1038/nrm3920

24 Csizmok, V., Follis, A. V., Kriwacki, R. W. & Forman-Kay, J. D. Dynamic Protein Interaction Networks and New Structural Paradigms in Signaling. Chem Rev 116, 6424–6462 (2016). 10.1021/acs.chemrev.5b00548

25 Sun, X., Rikkerink, E. H., Jones, W. T. & Uversky, V. N. Multifarious roles of intrinsic disorder in proteins illustrate its broad impact on plant biology. Plant Cell 25, 38–55 (2013). 10.1105/tpc.112.106062

26 Gao, Y., Li, X., Li, P. & Lin, Y. A brief guideline for studies of phase-separated biomolecular condensates. Nat Chem Biol 18, 1307–1318 (2022). 10.1038/s41589-022-01204-2

27 Wang, B. et al. Condensation of SEUSS promotes hyperosmotic stress tolerance in Arabidopsis. Nature Chemical Biology 18, 1361–1369 (2022). 10.1038/s41589-022-01196-z

28 Huang, X. et al. ROS regulated reversible protein phase separation synchronizes plant flowering. Nature Chemical Biology 17, 549–557 (2021). 10.1038/s41589-021-00739-0

29 Xie, Z. et al. Phenolic acid-induced phase separation and translation inhibition mediate plant interspecific competition. Nature Plants 9, 1481–1499 (2023). 10.1038/s41477-023-01499-6

30 Romero P, O. Z., Li X, Garner EC, Brown CJ, Dunker AK. Sequence complexity of disordered protein. Proteins (2001).

31 Yan, Z.-W. et al. Endocytosis-mediated entry of a caterpillar effector into plants is countered by Jasmonate. Nature Communications 14 (2023). 10.1038/s41467-023-42226-1

32 Stitz, M., Gase, K., Baldwin, I. T. & Gaquerel, E. Ectopic Expression of AtJMTinNicotiana attenuata: Creating a Metabolic Sink Has Tissue-Specific Consequences for the Jasmonate Metabolic Network and Silences Downstream Gene Expression Plant Physiology 157, 341–354 (2011). 10.1104/pp.111.178582

33 Park JH, H. R., Kim HB, Baldwin IT, Feldmann KA, Feyereisen R. A knock-out mutation in allene oxide synthase results in male sterility and defective wound signal transduction in Arabidopsis due to a block in jasmonic acid biosynthesis. Plant J. (2002).

34 Takaoka, Y. et al. A comprehensive in vitro fluorescence anisotropy assay system for screening ligands of the jasmonate COI1-JAZ co-receptor in plants. J Biol Chem 294, 5074–5081 (2019). 10.1074/jbc.RA118.006639

35 Huang H, W. C., Tian H, Sun Y, Xie D, Song S. Amino acid substitutions of GLY98, LEU245 and GLU543 in COI1 distinctively affect jasmonate-regulated male fertility in Arabidopsis. Sci China Life Sci. (2014). 10.1007/s11427-013-4590-1

36 Feys B, B. C., Penfold CN, Turner JG. Arabidopsis Mutants Selected for Resistance to the Phytotoxin Coronatine Are Male Sterile, Insensitive to Methyl Jasmonate, and Resistant to a Bacterial Pathogen. . Plant Cell. (1994).

37 Chico, J.-M. et al. Repression of Jasmonate-Dependent Defenses by Shade Involves Differential Regulation of Protein Stability of MYC Transcription Factors and Their JAZ Repressors in Arabidopsis. The Plant Cell 26, 1967–1980 (2014). 10.1105/tpc.114.125047

38 Oldfield, C. J. & Dunker, A. K. Intrinsically Disordered Proteins and Intrinsically Disordered Protein Regions. Annual Review of Biochemistry 83, 553–584 (2014). 10.1146/annurev-biochem-072711-164947

39 Feng, Z., Jia, B. & Zhang, M. Liquid–Liquid Phase Separation in Biology: Specific Stoichiometric Molecular Interactions vs Promiscuous Interactions Mediated by Disordered Sequences. Biochemistry 60, 2397–2406 (2021). 10.1021/acs.biochem.1c00376

40 Clough, S. J. & Bent, A. F. Floral dip: a simplified method forAgrobacterium-mediated transformation ofArabidopsis thaliana. The Plant Journal 16, 735–743 (2008). 10.1046/j.1365-313x.1998.00343.x

41 Jing, H. et al. Regulation of AUXIN RESPONSE FACTOR condensation and nucleo-cytoplasmic partitioning. Nature Communications 13 (2022). 10.1038/s41467-022-31628-2

42 Kosmacz, M. & Skirycz, A. The Isolation of Stress Granules From Plant Material. Current Protocols in Plant Biology 5 (2020). 10.1002/cppb.20118

43 Nguyen, S. H., Webb, H. K., Mahon, P. J., Crawford, R. J. & Ivanova, E. P. Natural insect and plant micro-/nanostructsured surfaces: an excellent selection of valuable templates with superhydrophobic and self-cleaning properties. Molecules 19, 13614–13630 (2014). 10.3390/molecules190913614

