## Supplementary Information for "Wounding perception by jasmonate-mediated JAZ condensation in plants"

**This PDF file includes:**

**Extended Data Fig.: 1 to 13**

**Description of Supplementary information:**

**Supplementary Table 1.** Primers used in this study.

**Supplementary Table 2.** Proteins identified from the JAZ1 condensates by LC-MS.

**Supplementary Video 1.** V-JAZ1 rapidly undergoes condensation after wounding. The pavement cells in Zone 1 were selected for laser wounding and imaging process was initiated immediately post wounding for a duration of 2 min. Time is shown at the bottom left. Scale bars, 5 $\mu\text{m}$.

**Supplementary Video 2.** Fusion events of V-JAZ1 condensates at Zone 2. Time is shown at the bottom left. Scale bar, 5 $\mu\text{m}$.

**Supplementary Video 3.** FRAP of JAZ1 condensates in Zone 2. The selected V-JAZ1 condensate was bleached and image was taken every three second and lasted for 1 min. Time is shown at the bottom left. Scale bar, 5 $\mu\text{m}$.

**Supplementary Video 4.** FRAP of JAZ1 condensates in *Arabidopsis* root cells of 35S:V-JAZ1 *Col-0* transgenic plants. The selected V-JAZ1 condensate was bleached and image was taken every three seconds and lasted for 1 min. Time is shown at the bottom left. Scale bar, 5 $\mu\text{m}$.

**Supplementary Video 5.** FRAP of JAZ1 condensates in *Arabidopsis* root cells of 35S:V-JAZ1 *jazQ* transgenic plants. The selected V-JAZ1 condensate was bleached and image was taken every three seconds and lasted for 1 min. Time is shown at the bottom left. Scale bar, 5 $\mu\text{m}$.

**Supplementary Video 6.** MeJA treatment triggers V-JAZ1 condensation. Image was taken every thirty seconds after MeJA addition for a duration of 20 min. Time is shown at the bottom left. Scale bar, 5 $\mu\text{m}$.

**Supplementary Video 7.** FRAP of mCherry-JAZ1 condensates *in vitro*. The selected mcherry-JAZ1 condensation was bleached and image was taken every one second and lasted for 1 min. Time is shown at the bottom left. Scale bar, 1 $\mu\text{m}$.

**Supplementary Video 8.** Fusion events of the mCherry-JAZ1 condensates *in vitro*. Image was taken every one second and lasted for 2 min. Time is shown at the bottom left. Scale bar, 1 $\mu\text{m}$.

**Supplementary Video 9.** JAZ1 undergoes condensation rapidly after MeJA treatment in *Arabidopsis* root cells. 35S:V-JAZ line 98 transgenic plants were treated with 50 $\mu\text{M}$ MeJA and the imaging process is started immediately after MeJA addition for a duration of 20 min. Time is shown at the bottom left. Scale bars, 5 $\mu\text{m}$.

**Supplementary Video 10.** JAZ1<sup>$\delta\text{ARR}$</sup> is dispersed in the nucleus of *Arabidopsis* root cells regardless of MeJA treatment. 35S:V-JAZ1<sup>$\delta\text{ARR}$</sup> line 32 transgenic plants were treated with 50 $\mu\text{M}$ MeJA, and the imaging process was started immediately after MeJA addition for a duration of 20 min.. Time is shown at the bottom left. Scale bars, 5 $\mu\text{m}$.

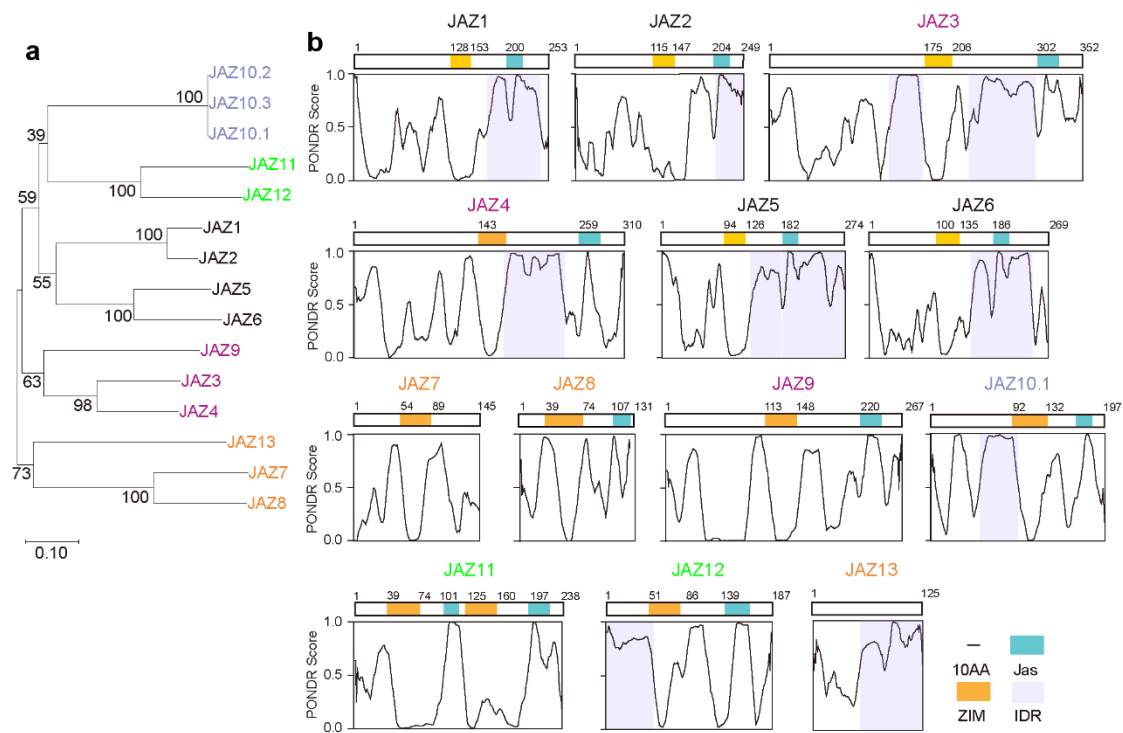

**Extended Data Fig. 1: Multiple JAZ proteins harbor IDRs.** **a**, Phylogenetic analysis of JAZ proteins. **b**, IDR distributions in JAZ protein sequences. The conserved domains of ZIM (orange) and Jas (blue) were marked in the top. Bottom is the prediction of the disordered regions (IDR, light purple) by the PONDR (www.pondr.com).

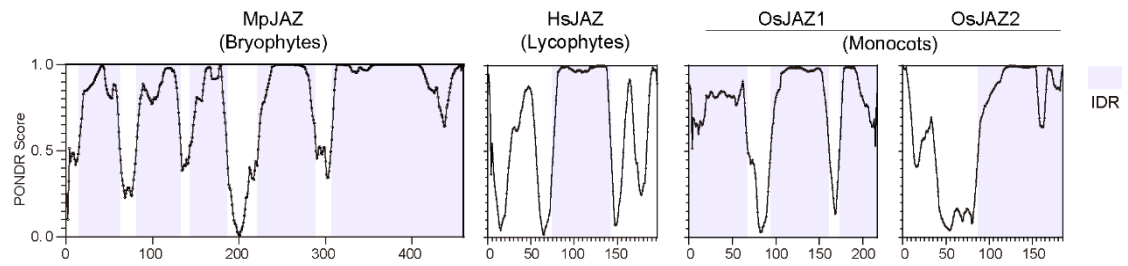

**Extended Data Fig. 2: JAZ proteins from multiple plant species harbor IDRs. IDR is marked in light purple.**

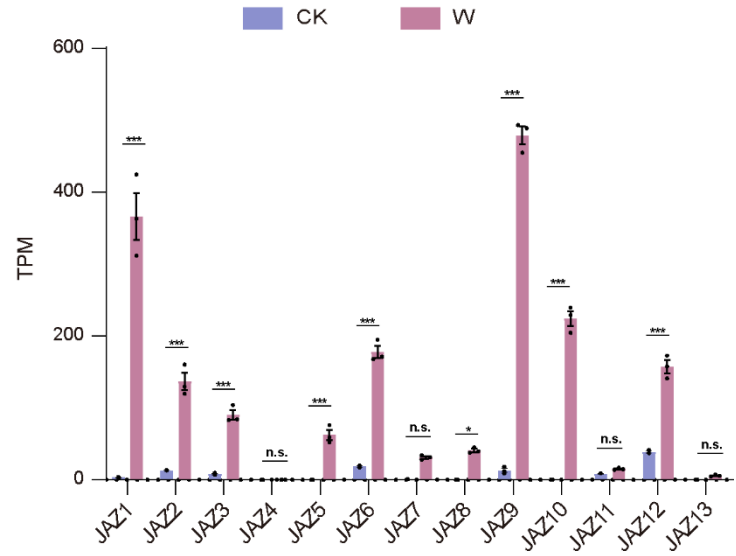

**Extended Data Fig. 3: The transcription level of JAZs from the RNA-seq data<sup>6</sup>.** CK represents unwounded plants (purple), and W represents wounding treated plants (red). Data are mean ± SEM (n = 3 biological replicates) and analyzed by two-sided Student's t-test. n.s.: no significance, \*$P < 0.05$, \*\*\*$P < 0.001$.

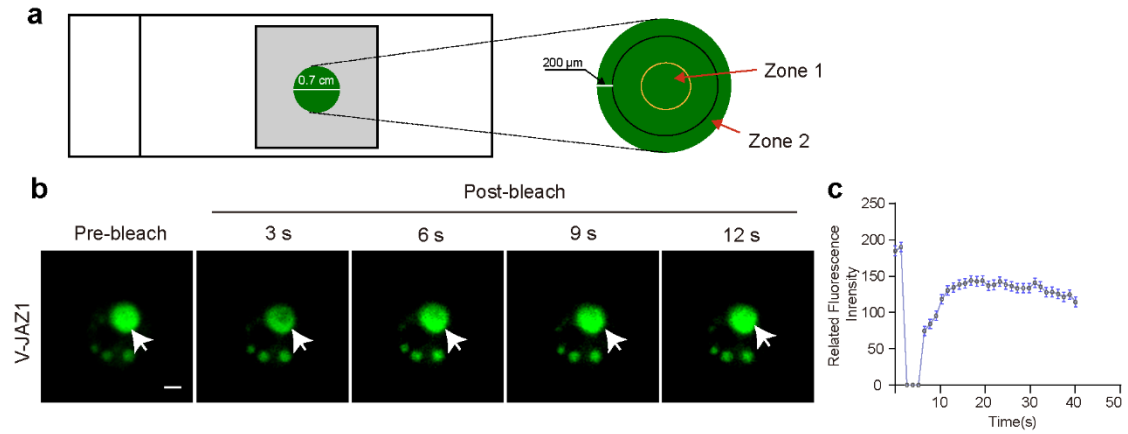

**Extended Data Fig. 4: JAZ1 undergoes liquid-liquid phase separation in *N. benthamiana*.**

**a**, Leaf disc used for microscopy observation. Zone 1 represents the center of the disc indicating the area far away from wounding side and Zone 2 represents the cut edge of the disc indicating the area around the wounding site. **b**, Fluorescence recovery after photobleaching (FRAP) at Zone 2. The white arrow indicates the selected condensate for photobleaching. Scale bars, 5 μm. **c**, Quantification of the recovered fluorescence intensity of the condensates as described in (b). Data are mean ± SEM (n = 6 biological replicates).

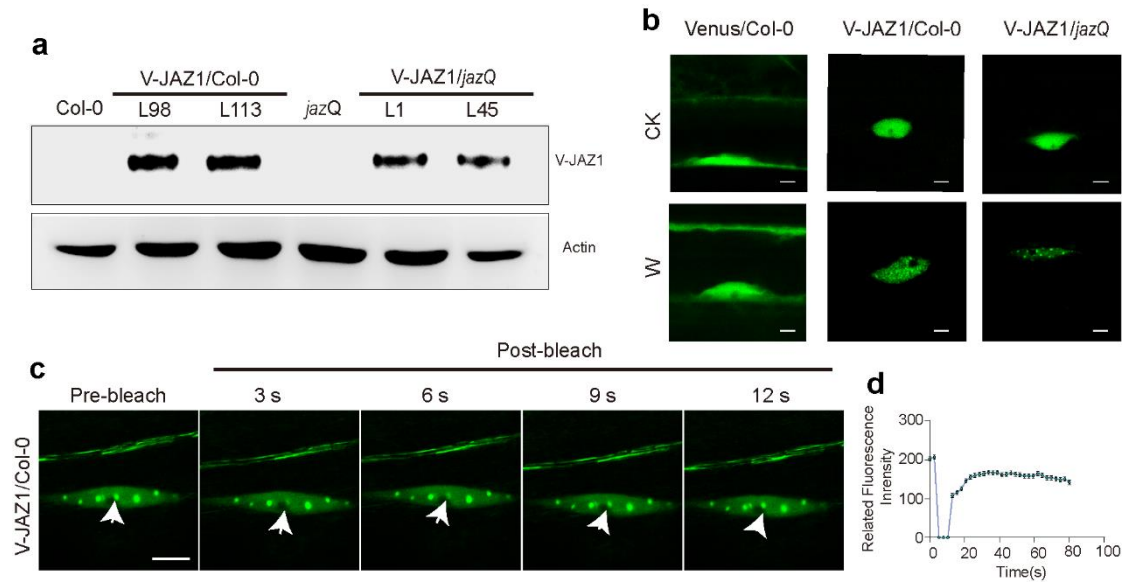

**Extended Data Fig. 5: JAZ1 undergoes liquid-liquid phase separation in *Arabidopsis*.** **a**, The protein levels of V-JAZ1 in Col-0, *jazQ* and transgenic plants. Anti-Venus antibody is used to detect V-JAZ1. Anti-Actin antibody is used to detect Actin. **b**, JAZ1 forms condensates in both 35S:*V-JAZ1* line 113# (V-JAZ1/Col-0) and 35S:*V-JAZ1 jazQ* line 45# (V-JAZ1/*jazQ*) upon wounding. Venus-expressed plants (35S:*Venus*, Venus/Col-0) were used as parallel control. The untreated (CK) and wounded (W) cells of the root elongation region are observed under confocal microscopy. Scale bars, 5 μm. **c**, FRAP analysis of JAZ1 condensates in root cells of 35S:*V-JAZ1* line 98# (V-JAZ1/Col-0). The white arrows indicate the selected condensate for photobleaching. Scale bars, 5 μm. **d**, Quantification of the recovered fluorescence intensity of the condensates as described in (c). Data are mean ± SEM (n = 6 biological replicates).

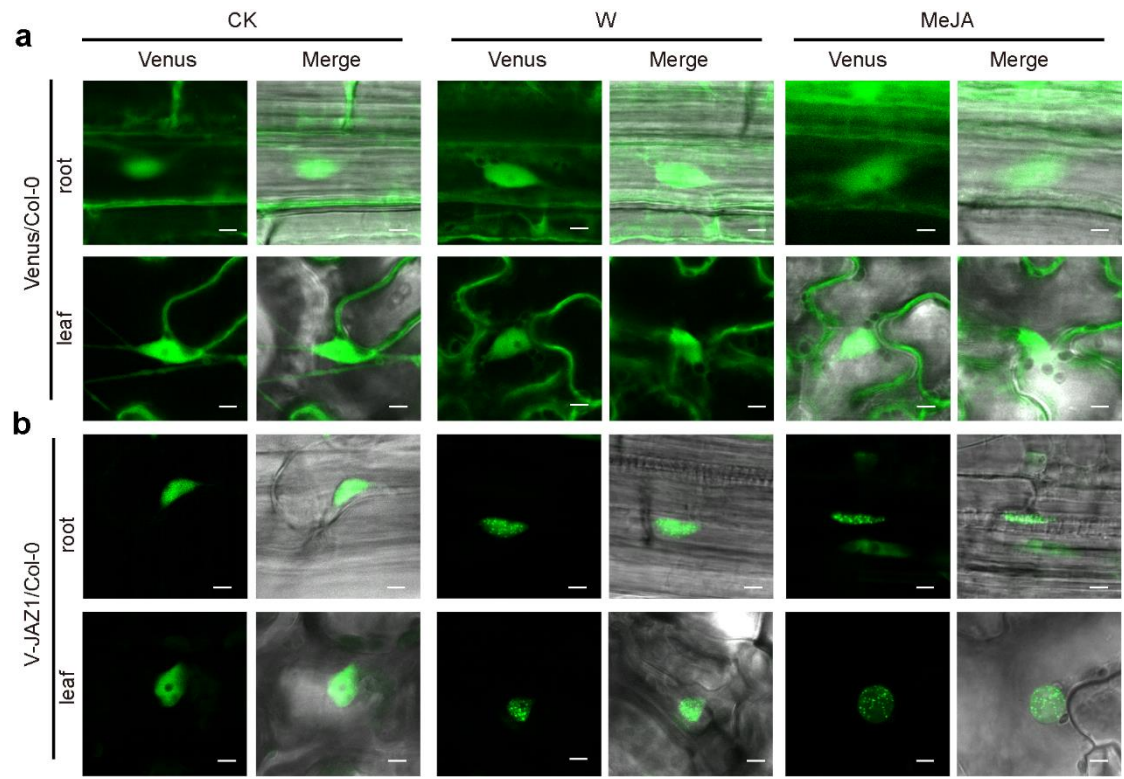

**Extended Data Fig. 6: The condensation of JAZ1 is triggered by both wounding and MeJA treatments.** The plants of 35S:*Venus* (Venus/Col-0) and 35S:V-JAZ1 line113 (V-JAZ1/Col-0) were wounded or treated with 50 μM MeJA. The untreated (CK), wounding treated (W) and MeJA treated (MeJA) plants were observed under confocal microscopy. **a**, Venus is unable to form condensates. **b**, JAZ1 undergoes condensation in plant response to both wounding and MeJA. Scale bars, 5 μm.

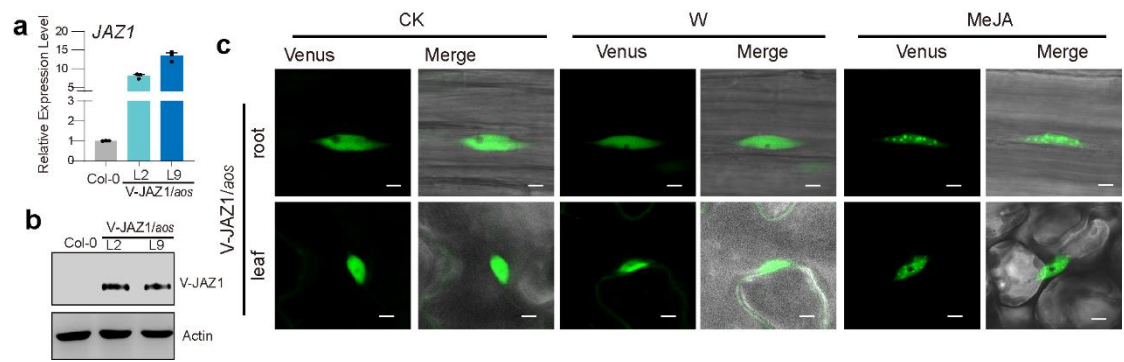

**Extended Data Fig. 7: Wounding-mediated JAZ1 condensation is dependent on JA.** **a**, qRT analysis of *JAZ1* expression in Col-0 and *35S::V-JAZ1 aos* plants. Data are mean ± SEM ($n = 3$ biological replicates). **b**, Immune blot assay of V-JAZ1. Anti-Venus antibody is used to detect V-JAZ1. Anti-Actin antibody is used to detect Actin. **c**, MeJA but not wounding treatment triggers JAZ1 condensation in *35S::V-JAZ1 aos* line 9 (V-JAZ1/*aos*). Plants were treated with wounding or 50 μM MeJA. The root and leaf cells of the untreated (CK), wounded (W) and MeJA-treated (MeJA) plants were observed under confocal microscopy. Scale bars, 5 μm.

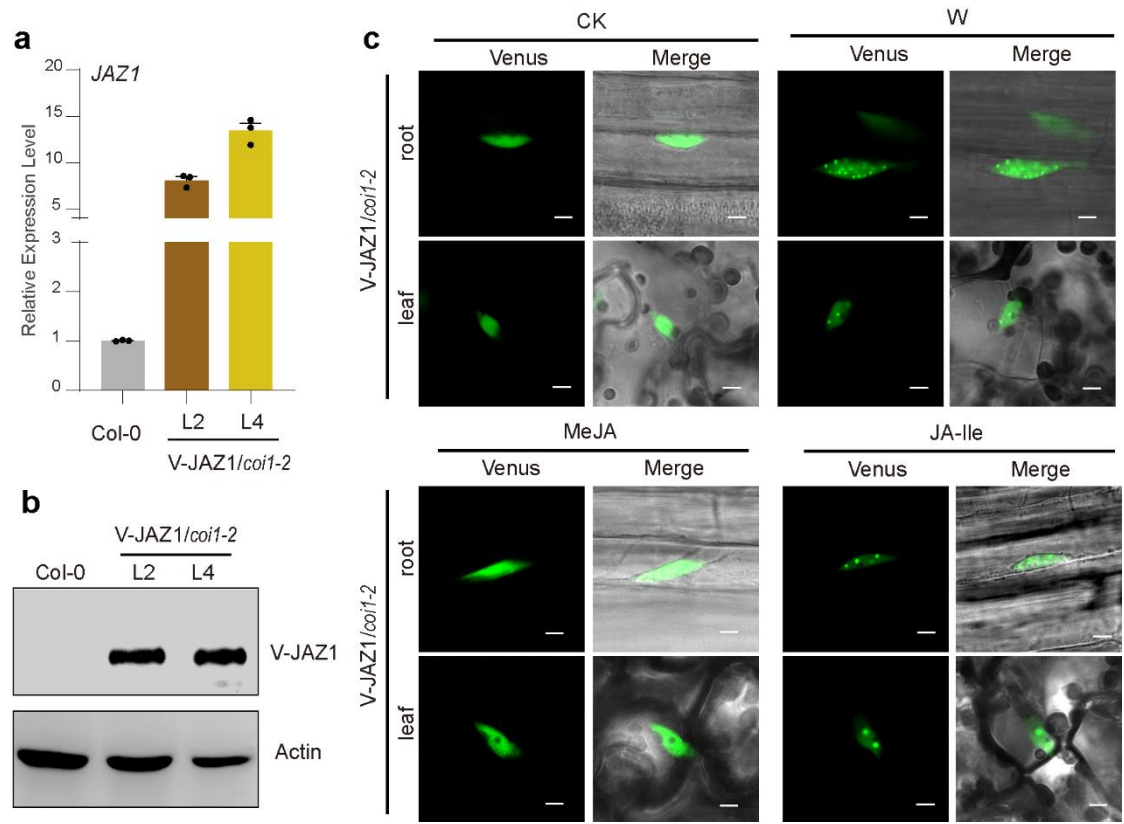

**Extended Data Fig. 8: JAZ1 condensation is independent on COI1.** **a**, qRT analysis of *JAZ1* expression in Col-0 and *35S:V-JAZ1 coi1-2* plants. Data are mean ± SEM ($n = 3$ biological replicates). **b**, Immune blot assay of JAZ1. Anti-Venus antibody is used to detect V-JAZ1. Anti-Actin antibody is used to detect Actin. **c**, JAZ1 condensation was observed in response to wounding and JA-Ile treatments in *35S:V-JAZ1 coi1-2* plants (*V-JAZ1/coi1-2*). The plants of line 4# were wounded (W) or treated with 50 μM of MeJA and JA-Ile respectively. The untreated plants (CK) and plants with indicated treatment were observed under confocal microscopy. Scale bars, 5 μm.

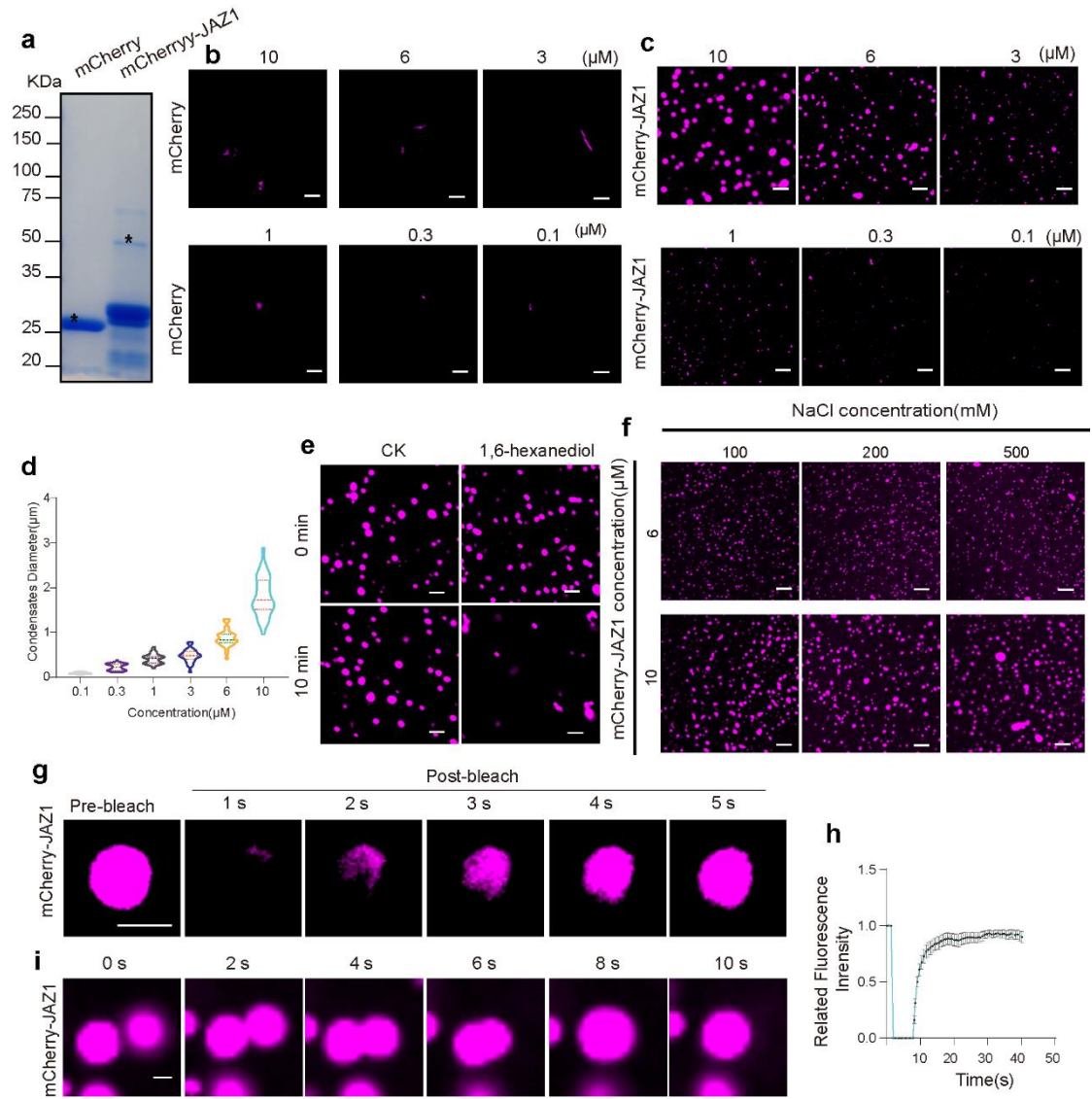

**Extended Data Fig. 9: JAZ1 undergoes liquid-liquid phase separation *in vitro*.** **a**, Coomassie blue staining. The mCherry and mCherry-fused JAZ1 (mCherry-JAZ1) protein were expressed in prokaryotes and purified for SDS-PAGE analysis. mCherry and mCherry-fused JAZ1 were marked with a star. **b-c**, The purified mCherry (b) and mCherry-JAZ1 (c) of indicated concentrations were observed under confocal microscopy. Scale bars, 5 $\mu\text{m}$. **d**, Diameter measurement of the condensates described in (c). Data are mean ± SEM (n = 35). **e**, 1,6-hexanediol disrupts mCherry-JAZ1 condensation. 10 $\mu\text{M}$ mCherry-JAZ1 solutions are incubated without (CK) or with 10% 1,6-hexanediol for 10 min. Scale bars, 5 $\mu\text{m}$. **f**, The presence of NaCl has little effect on mCherry-JAZ1 condensates. The mCherry-JAZ1 solutions of 6 $\mu\text{M}$ and 10 $\mu\text{M}$ were mixed with NaCl of indicated concentrations for 10 min before being used for confocal observation. Scale bars, 5 $\mu\text{m}$. **g**,

114 FRAP assay of mCherry-JAZ1 condensates. Scale bars, 1 $\mu\text{m}$. **h**, Quantification of the
115 recovered fluorescence intensity after photobleaching described in (g). Data are mean ± SEM
116 (n = 6 biological replicates). **i**, The mCherry-JAZ1 condensates are fusing together. Scale bar,
117 1 $\mu\text{m}$.

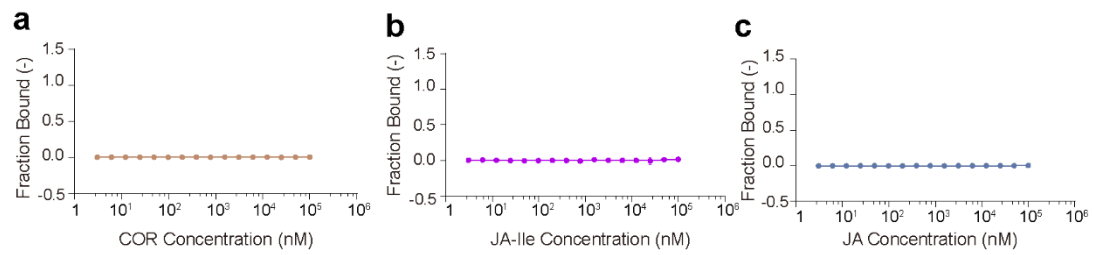

**Extended Data Fig. 10:** MST assay shows that mCherry do not bind with the Coronatine (COR), JA-Ile and JA, Yellow purple and green line represent COR (a), JA-Ile (b) and JA (c). Data are mean ± SD (n = 3 biological replicates).

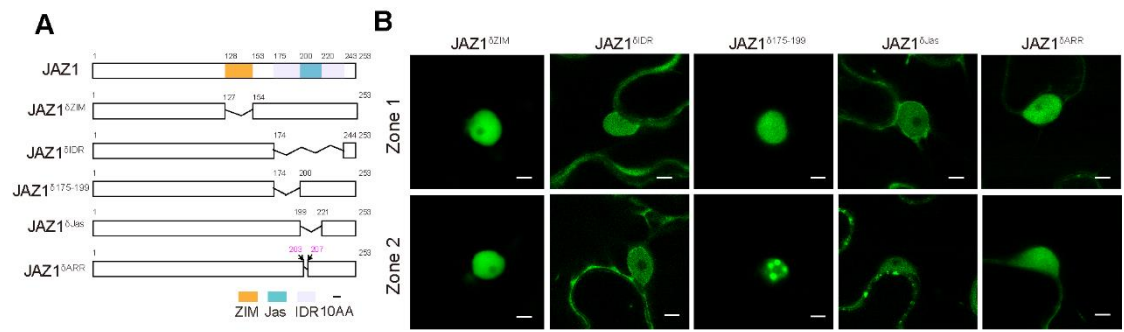

**Extended Data Fig. 11: Condensation of the truncated JAZ1 variants in JA response. a,** Schematic of JAZ1 protein sequence and its truncated variants. Orange, blue and light purple boxes represent the ZIM, Jas domain and IDR respectively. Lines indicate mutated positions. **b,** Observation of truncated V-JAZ1 variants at Zone1 and Zone2. The truncated V-JAZ1 variants were transiently expressed in the leaves of *N. benthamiana*, and observed under confocal microscopy. Scale bars, 5 μm.

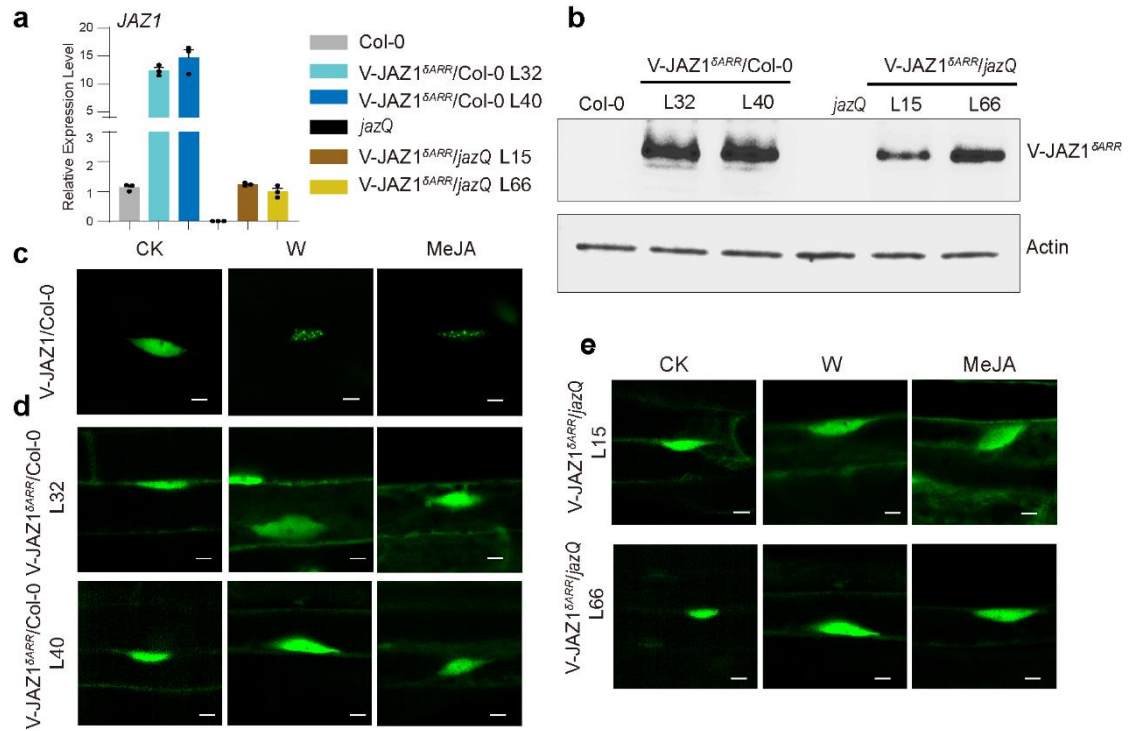

**Extended Data Fig. 12: JAZ1<sup>δARR</sup> fails to form condensates in response to wounding and MeJA treatment in *Arabidopsis*.** **a** qRT analysis of *JAZ1* in Col-0, *jazQ* and transgenic plant. Data are mean ± SEM (n = 3 biological replicates). **b**, Immune blot assay of V-JAZ1<sup>δARR</sup>. Anti-Venus antibody is used to detect V-JAZ1<sup>δARR</sup>. Anti-Actin antibody is used to detect Actin. **c-e**, Observations of V-JAZ1 and V-JAZ1<sup>δARR</sup> in *Arabidopsis*. Plants of 35S:*V-JAZ1* line 99# (V-JAZ1/Col-0), 35S:*V-JAZ1*<sup>δARR</sup> line 32 (V-JAZ1<sup>δARR</sup>/Col-0 L32), 35S:*V-JAZ1*<sup>δARR</sup> line 40 (V-JAZ1<sup>δARR</sup>/Col-0 L40), 35S:*V-JAZ1*<sup>δARR</sup> *jazQ* line 15 (V-JAZ1<sup>δARR</sup>/*jazQ* L15) and 35S:*V-JAZ1*<sup>δARR</sup> *jazQ* line 66 (V-JAZ1<sup>δARR</sup>/*jazQ* L66) were wounded or treated with 50 μM MeJA. The untreated (CK), wounded (W) and MeJA-treated (MeJA) plants were observed under confocal microscopy. Scale bars, 5 μm.

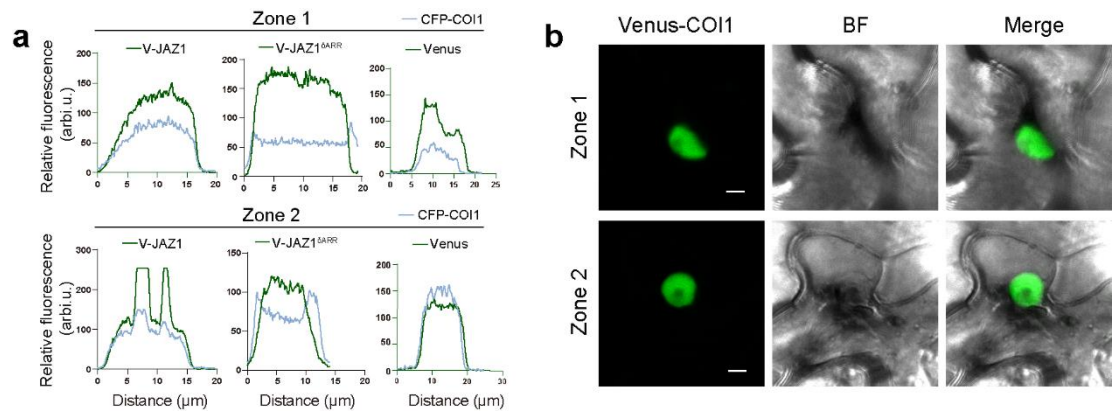

**Extended Data Fig. 13: COI1 is failed to form condensates in Zone 2.** **a**, Co-localization of CFP-COI1 and V-JAZ1 as described in Fig. 4d. Fluorescence intensity (in arbitrary units, arbi.u.) in cross-section was quantified. CFP-COI1 was transiently co-expressed with V-JAZ1 and Venus in leaves of *N. benthamiana*, respectively. The green lines indicate the intensity of V-JAZ1 or Venus. The blue lines indicate the intensity of CFP-COI1. **b**, No COI1 condensate was observed in both Zone1 and Zone 2. The Venus fused COI1 (Venus-COI1) was transiently expressed in *N. benthamiana* leaves and observed under microscopy. Scale bar, 5 μm.

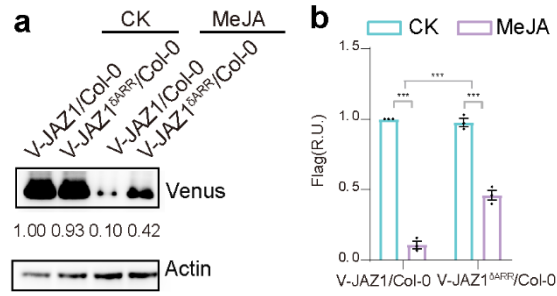

**Extended Data Fig. 14: JAZ1 is more easily degraded than V-JAZ1<sup>ΔARR</sup> in response to MeJA.** **a**, V-JAZ1 and V-JAZ1<sup>ΔARR</sup> levels in plants by immune blot assay. *35S::V-JAZ1* line 98 (V-JAZ1/Col-0) and *35S::V-JAZ1<sup>ΔARR</sup>* line 32 (V-JAZ1/Col-0) were treated by 50 μM MeJA and collected 1 hour post treatment. V-JAZ1 and V-JAZ1<sup>ΔARR</sup> in the untreated (CK) and MeJA-treated (MeJA) plants were detected. Anti-Venus antibody was used to detect V-JAZ1 and V-JAZ1<sup>ΔARR</sup>. Anti-Actin antibody is to detect Actin proteins as loading controls. **b**, Quantification of V-JAZ1 and V-JAZ1<sup>ΔARR</sup> as described in (a). Data are analyzed by two-way ANOVA followed by multiple comparisons with two-sided Fisher's LSD test (\*\*\*) $P < 0.001$. Data are mean ± SEM (n = 3 biological replicates).
